# Impaired Reinforcement Learning Underlying Explore-Exploit Decision Making in Theft Recidivists

**DOI:** 10.64898/2026.09.11.750836

**Authors:** Yukiori Goto, Satoshi Yoshino, Chikara Kita, Moojun Won, Young-A Lee

**Author notes:** **Address Correspondence:** Yukiori Goto, Ph.D., Department of Artificial Intelligence and Technology Graduate School of Informatics, Kyoto University, Yoshida-hanamachi, Sakyo-ku, Kyoto 606-8501, Japan.

## Abstract

Larceny imposes profound societal and economic burdens; however, punitive judicial measures frequently fail to deter recidivism. The neurobehavioral mechanisms driving habitual offending, whether instrumental or kleptomanic, in theft recidivists remain poorly understood. In this study, we investigated explore-exploit decision-making and underlying reinforcement learning architectures in theft recidivists with a 4-arm bandit task while prefrontal cortex (PFC) hemodynamics were continuously monitored using functional near-infrared spectroscopy (fNIRS). Model-free behavioral analyses revealed that non-kleptomanic (TR-K) but not kleptomanic (TR+K) theft recidivists accumulated significantly higher cumulative regret and made fewer optimal choices compared to control individuals without criminal records (CT). Model-based analyses through Variational Bayesian Analysis identified Q-learning with decay model as the optimal computational fit. Parameter extraction using the model demonstrated that the TR-K group exhibited a lower learning rate than CT and TR+K groups, indicating a learning deficit in updating action values following environmental feedback. fNIRS tracking of trial-by-trial latent reinforcement variables revealed that while PFC activity was modulated by these variables, group differences were characterized by static baseline hemodynamic shifts rather than rewirings of value-tracking neural circuits. These findings suggest that non-kleptomanic recurrent thefts may be associated with impaired reinforcement learning mechanisms, challenging current punitive deterrence models.

**SIGNIFICANCE STATEMENT:** This study shows a neurobehavioral mechanism underlying recurrent theft, challenging the traditional assumption that recidivism stems from rational choice or moral failure. By utilizing a reinforcement learning framework, this study demonstrates that instrumental, but not kleptomanic, theft recidivists possess a diminished learning rate. Such learning deficit prevents them from efficiently updating the expected value of their actions following feedback, such as incarceration. Consequently, standard punitive deterrence models fail for this specific population. These findings highlight the need to pivot justice systems toward neurobehavioral rehabilitation strategies that target reinforcement learning deficits.

## INTRODUCTION

Larceny, particularly shoplifting, imposes profound societal and economic burdens globally (Blanco et al., 2008; Yaniv, 2009). Despite judicial punitive measures, including incarceration, the rate of recidivism among theft offenders remains high (Fazel & Wolf, 2015; Yukhnenko et al., 2023). Although the criminal justice system implicitly assumes that offenders possess the cognitive architecture required to adjust their future behavior in response to severe negative consequences (Becker, 1968), the chronic nature of recurrent theft suggests a fundamental breakdown in this adaptive process. Thus, repeated offending, particularly in the case of psychopathy, may be associated with less of a calculated choice and more reflective of underlying cognitive and emotional processing associated reinforcement learning with a persistent failure to learn from negative environmental feedback (Blair et al., 2004; Hosking et al., 2017).

Accumulating evidence suggests that theft recidivists constitute a clinically or neurocognitively heterogeneous population (Goto et al., 2026; Grant, 2006). In particular, a clinical distinction exists between instrumental theft recidivists, who commit theft driven by profit, socioeconomic necessity, or broader antisocial tendencies, and those diagnosed with kleptomania, an impulse-control disorder characterized by recurrent, irresistible urges to steal items with little to no monetary value (Association, 2013). While the psychiatric and neurobiological underpinnings of kleptomania, including its addictive-like compulsions, have received attention (Asaoka et al., 2025; Grant, 2006), a critical knowledge gap remains.

Computational psychiatry offers a robust theoretical framework for reinforcement learning (RL) (Adams et al., 2016; Maia & Frank, 2011; Montague et al., 2012). RL models posit that individuals navigate their environments by continuously updating the expected value, or Q-value, of available actions based on environmental feedback. This value-updating process is driven by the reward prediction error (RPE), which is the quantifiable discrepancy between an expected and actual received outcomes (Lowet et al., 2020; Schultz et al., 1997; Sutton & Andrew, 2018). A fundamental challenge in RL is the explore-exploit dilemma: the cognitive demand to balance exploitation that involves choosing a known, safe option to maximize a predictable immediate reward based on current knowledge against exploration that involves choosing an unknown or previously low-value option to seek novel information or potentially higher future rewards (Addicott et al., 2017; Cohen et al., 2007). In particular, under this framework, instrumental theft may be considered as a salient exploitative behavior, whereas kleptomania may represent a dysregulation of exploration and exploitation balance rather than rational exploitation.

The multi-armed bandit task has emerged as a psychological paradigm to assess RL dynamics (D. et al., 2017; Daw et al., 2006). Studies utilizing the bandit task have demonstrated that healthy individuals seamlessly alternate between exploitation and exploration, a process guided by individual learning rates and decision temperatures (Seymour et al., 2012; Speekenbrink & Konstantinidis, 2015), whereas in psychiatric populations, such as those with pathological gambling and substance use disorder, aberrant exploit-explore decision making, often manifesting as learning rigidity, has been reported (Harle et al., 2015; Hodson et al., 2026; Wiehler et al., 2021). The prefrontal cortex (PFC) has been shown to play a significant role in these RL computations. In particular, the dorsolateral PFC maintains task rules, overrides prepotent responses, and tracks specific action values, whereas the dorsomedial PFC calculates RPEs, evaluates the consequences of actions, and triggers subsequent behavioral shifts (Barraclough et al., 2004; Domenech et al., 2020; Rushworth et al., 2011; Sazhin et al., 2025; Seo et al., 2007).

This study aimed to investigate behavioral and neural differences in explore-and-exploit decision-making and relevant reward learning across three distinct populations: healthy controls with no history of criminal records (CT), theft recidivists diagnosed with kleptomania (TR+K), and theft recidivists without kleptomania (TR-K). We employed a 4-arm bandit task integrated with Bayesian computational modeling to extract individual reinforcement learning parameters. Concurrently, we utilized functional near-infrared spectroscopy (fNIRS) to sample PFC responses during task execution.

## METHODS

### Subjects

Sixty-three theft recidivists (TR) and 53 control subjects (CT) were recruited for this study, along with our other studies. The inclusion criteria for the TR group were prior records of incarceration due to larceny (primarily shoplifting), being between 18 and 79 years of age, and living in Japan at the time of the study. The CT group inclusion criteria required participants to have no prior criminal records, be between 18 and 79 years of age, and reside in Japan. The exclusion criteria for all participants was the inability to understand the study details due to intellectual disability or any other cognitive impairment, for which no recruited participants met these exclusion criteria. Socioeconomic status was not considered for inclusion or exclusion criteria, as all recruited TR participants were residing in rehabilitation centers or welfare facilities for societal reintegration, rendering their socioeconomic statuses relatively invariable.

The TR group was divided into 16 participants who had been formally diagnosed with kleptomania and were receiving clinical treatment at the time of the study (TR+K), and 47 participants who had no clinical diagnosis of kleptomania (TR-K). Prior to or during the task, 4 TR+K and 9 TR-K participants dropped out of the study due to a loss of interest. Consequently, the final sample size for behavioral analysis was conducted with 53 CT, 12 TR+K, and 38 TR-K participants.

This study was conducted in accordance with the Declaration of Helsinki and the Ethical Guidelines for Medical and Health Research Involving Human Subjects of the Japanese Ministry of Health, Labour, and Welfare. All procedures were approved by the Human Research Ethics Committee of the Kyoto University Graduate School of Informatics. Written informed consent was obtained from all participants prior to their involvement. Demographic variables, including age, sex, and smoking status, were recorded at the time of enrollment.

### 4-arm Bandit Task

To understand decision-making under uncertainty, a 4-arm bandit task, where payouts drift over time, was used. The task was programmed and administered to participants using Inquisit software (Millisecond Software, LLC., Seattle, WA, USA). In this task, a participant must repeatedly choose between four options (“arms”), each yielding rewards (payoffs) with unknown probabilities. The goal is to maximize the total reward by balancing exploration (trying different arms to learn their payouts) and exploitation (choosing the known best arm).

The task consisted of 160 trials, divided into two 80-trial blocks with a brief 15-second resting period. This number of trials was shorter than the originally used in other studies (Daw et al., 2006); however, this shortened version of the task was utilized because initial piloting with larger trial numbers (e.g., 300 trials) resulted in a high dropout rate, particularly in the TR group. Five demo trials were administered for practice before starting the test. At the beginning of each selection trial, 4 arms represented by different colors were presented on the screen. A different starting payoff value (initial means of 20, 40, 60, 80) was assigned to each slot, and these payoff values were constantly updated from trial to trial by incorporating a decaying Gaussian random walk for payoff distributions (enforced range 1-100).

Participants were explicitly instructed to maximize their payoffs by choosing one of 4 arms that was assumed to be the highest payoff. In addition, they were instructed that they had 1500 ms to make an arm selection; otherwise, they would lose the opportunity to gain a payoff in the trial. Accordingly, the trial timeline started at arm selection (maximum 1500 ms), followed by an animated slot (2000 ms), subsequently revealing the payoff (1000 ms). Subsequently, a blank screen appeared before starting the next trial (1000 ms). If a participant failed to make a selection within the 1500 ms window, an error feedback screen was displayed for 4000 ms, followed by a 1000 ms blank screen.

Both model-free and model-based measurements were evaluated in this task. Model-free behavioral metrics are raw, observable data to describe decision-making behavior, including cumulative regret (the difference between the maximum possible reward and the actual reward obtained), optimal choice proportion (the percentage of trials where the objectively highest-paying arm was selected), exploitative choice proportion (the percentage of trials that a participant selected the slot with the highest known payoff at this point), switch rate (the overall frequency of changing arms, serving as a raw proxy for exploratory behavior), and win-stay/lose-shift rates (the probability that a participant repeats a choice after a higher reward and switching after a lower reward than expected), which were calculated using the mathematical definitions in Suppl. Methods.

Model-based parameters are latent cognitive parameters extracted from RL model fitting to trial-by-trial data, including the estimated learning rate α (the measurement for updates of belief about an arm’s value after receiving a reward), inverse temperature β (the measurement of the trade-off between exploration and exploitation derived from Softmax decision models), decay rate γ (the measurement of regression for the estimates of unchosen arms over time), action value Q (the measurement for updates of the estimate of what each of the four arms is currently worth), and RPE (the measurement for the difference between the actual and expected payoff to receive from the chosen arm), which were derived as described in the data analysis below.

## fNIRS

To investigate the neural correlates of the RL process, fNIRS sampling was conducted. Given the environmental and logistical constraints for the TR+K group, sampling was conducted only in CT and TR-K participants. Measurements were obtained using the NIRO-200 NIRS Image Processing and Measuring System (Hamamatsu Photonics K.K., Hamamatsu, Japan). The system utilized two emitters delivering laser pulses at wavelengths of 775, 810, and 850 nm, and eight detectors with a 3.0 cm source-detector separation. Changes in oxygenated (O2Hb) and deoxygenated (HHb) hemoglobin were sampled at a rate of 0.5 Hz across 10 locations (R1‒R10) spanning the left and right hemispheres of the PFC.

Based on the MarsAtlas (Auzias et al., 2016), the channel locations were mapped as follows: R4 and R6 covered the left and right rostral dorsal PFC (PFrd; Brodmann area [BA] 10/9/8); R5 and R7 covered the left and right caudal dorsomedial PFC (PFcdm; BA 6/8); R3 and R8 covered the left and right rostral dorsolateral superior PFC (PFrdls; BA 10/9); R2 and R9 covered the left and right caudal dorsolateral PFC (PFcdl; BA 45/46/9); and R1 and R10 covered the intermediate zones between the PFrdls and PFcdl.

Since each trial of the 4-arm Bandit Task lasted a minimum of 4000 ms plus the reaction time (up to 1500 ms), fNIRS yielded two to three samples per trial by the sampling rate at 0.5 Hz. These samples were standardized, averaged for each trial, and subsequently utilized in data analysis.

### Data Analysis

All statistical and computational analyses were conducted using JASP ver. 0.97.1 (Team, 2026), OriginPro ver. 2026 (OriginLab Corporation, Northampton, MA, USA), and MATLAB R2026a (The MathWorks Inc., Natick, MA, USA). Bayesian statistical analysis was employed throughout the study, except for model validations, as it provides a robust alternative to frequentist methods when dealing with relatively small and severely unbalanced sample sizes between groups.

#### Model-free measurements

Bayesian ANCOVA was conducted to evaluate the effect of Group (CT vs. TR+K vs. TR-K) on behavioral variables including cumulative regret, proportion of optimal choice, proportion of exploitative choice, switch rate, corrected win-stay rate, and corrected lose-shift rate, while statistically controlling for Age (18-78), Sex (Male vs. Female), and Smoking status (Yes vs. No). In particular, a main-effects model was compared with a null hypothesis with covariates (covariate-only) model. Default prior distributions were employed for the analysis. Thus, a multivariate Cauchy (Jeffreys-Zellner-Siow; JZS) prior was placed on the standardized effect sizes, with a default scale parameter of r = 0.5 for fixed effects and r = 0.354 for covariates (Rouder & Morey, 2012).

The relative evidence for each model was quantified using the Bayes Factor (BF10), which calculates the probability of the observed data under a given model relative to the null model. A BF10 >1 indicates evidence in favor of the alternative model, whereas BF10 <1 indicates evidence for the null. The strength of the evidence was interpreted according to the classification scheme provided by Lee and Wagenmakers (2013) (Lee & Wagenmakers, 2014): 1 < BF10 < 3 suggests anecdotal evidence, 3 < BF10 < 10 suggests moderate evidence, 10 < BF10 < 30 10 suggests strong evidence, and BF10 > 30 suggests very strong to extreme evidence.

#### Model-based measurements

Computational model inversion was performed using the Variational Bayesian Analysis (VBA) toolbox (Daunizeau et al., 2014) in MATLAB, which is the toolkit for fitting non-linear dynamical systems to behavioral choice data. This requires mapping the RL process into a state-space model by defining how the agent’s internal values update (the evolution function) and how those values translate into actual choices (the observation function). We fitted three RL models to the behavioral data: a standard Q-learning model using a Softmax observation rule, a Q-learning model with a decay parameter for unchosen options, and a random choice model (Suppl. Methods). The evolution and observation parameters were estimated by maximizing the variational Free Energy under Laplace approximation.

To evaluate model fitting, the Bayesian Information Criterion (BIC) and McFadden’s pseudo-R^2^ calculated as 1-(lnLmodel/lnLrandom), which represented the improvement in model fit over the random choice baseline, were computed for each subject (Suppl. Table S1). To determine the computational architecture that best described subject behavior, group-level Bayesian Model Selection (BMS) was performed on the model evidence (Free Energy) to compute the exceedance probability for each model, which measured the likelihood that a given model was more frequent in the population than any other model, penalized for model complexity. Following model selection, subject-specific maximum a posteriori (MAP) parameters (learning rate α, inverse temperature β, and decay rate γ) of the winning model were extracted. To evaluate group differences while controlling relevant covariates, a Bayesian ANCOVA was conducted on these parameters across the CT, TR+K, and TR-K groups.

To ensure the robustness of modeling, we also conducted a cross-fitting model recovery analysis to evaluate whether the BMS procedure could reliably distinguish between standard Q-learning and Q-learning with decay models, as well as a parameter recovery analysis to confirm that the parameters (α, β, and γ) of the winning model were independently identifiable (Suppl. Methods). In addition, to validate the behavioral predictive power of the model, predictive choice probabilities and trial-by-trial Q-values were reconstructed for individual subjects. Learning curves were generated by mapping the pre-update Q-values against actual participant choices (weighted by normalized reward outcomes 0-1). Model accuracy was evaluated by calculating the percentage of trials in which the highest predicted choice probability of the model matched the empirical action, confirming that the latent state trajectories meaningfully captured human behavioral dynamics.

#### Bayesian Linear Mixed-effect Model (LMM) for fNIRS Analysis

Two trial-by-trial latent reinforcement variables, Q-value and RPE, were extracted from the winning computational model fitting to the behavioral data. To evaluate the neural correlates of these RL variables and assess between-group differences, data were analyzed using Bayesian LMMs with the Q-value and RPE used as predictors for the fNIRS signals. Thus, models evaluated the fixed effects of Group (TR-K vs. CT), Q-value, RPE, and their interactions (Group x Q-value; Group x RPE) on trial-by-trial standardized O2Hb and HHb fNIRS signals across the 10 channels. Subject-level variations were accounted for via random intercepts.

To ensure robust estimation of the posterior distributions, the Markov chain Monte Carlo (MCMC) sampling was configured with 4 chains. Each chain was run for 8,000 iterations after a burn-in period of 4,000 iterations. To optimize sampling efficiency and avoid divergent transitions in the complex multidimensional posterior space, the sampler settings were tightly controlled with an adapt delta of 0.95 and a maximum treedepth of 12. The statistical credibility of the fixed effects and contrasts was determined using 95% High Posterior Density Intervals (HPDI), such that an effect was considered credible if its 95% interval did not contain zero.

## RESULTS

### Model-free measurements

Model-free behavioral metrics were analyzed using Bayesian ANCOVA. The primary model evaluated the effect of Group (CT [n=53], TR+K [n=12], and TR-K [n=38]) against a null hypothesis with covariates model, with Age, Sex, and Smoking status included as covariates.

The analysis revealed decisive evidence for a group effect on cumulative regret (BF10 = 463.4). Post-hoc comparisons demonstrated decisive evidence for a difference between the CT and TR-K groups (BF10 = 425.9; Fig. 1a; Suppl. Table S1), while comparisons between other groups did not exceed a Bayes factor of 1. Similarly, strong evidence for a group effect was observed for the proportion of optimal choice (BF10 = 39.06; post-hoc CT vs. TR-K: BF10 = 24.89; Fig. 1b; Suppl. Table S1) and the proportion of exploitative choice (BF10 = 54.36; post-hoc CT vs. TR-K: BF10 = 25.19; Fig. 1c; Suppl. Table S1).

**Figure 1.**
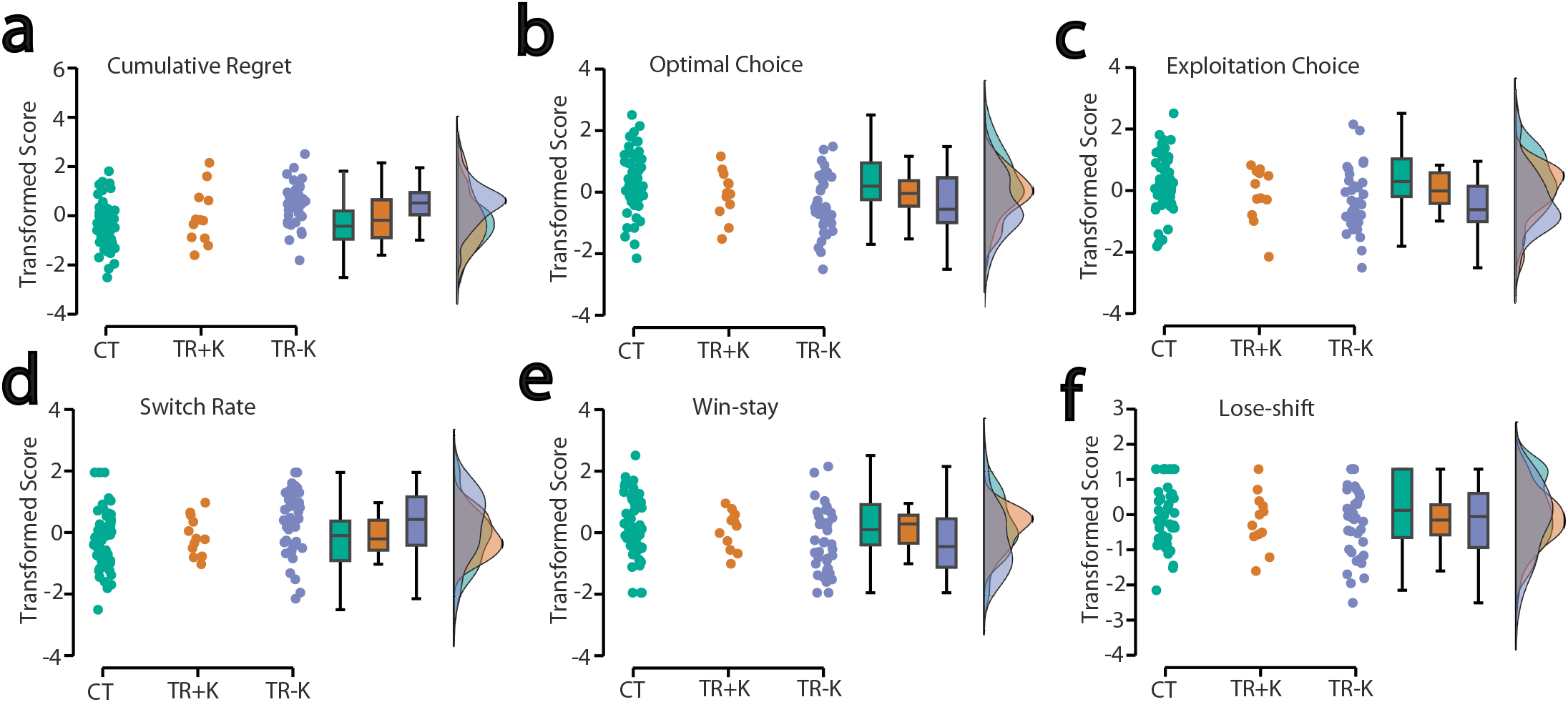
Model-free task measurements in TR+K, TR-K, and CT. a-f, Raincloud plots comparing transformed (data with rank-based inverse normal transformation) scores of empirical cumulative regret (a), proportion of optimal choice (b), proportion of exploitative choice (c), switch rate (d), chance-corrected win-stay rate (e), and chance-corrected lose-shift rate between theft recidivists with kleptomania (TR+K), those without kleptomania (TR-K), and participants with no criminal records (CT).

We also found moderate evidence for group differences in behavioral flexibility measures. The switch rate showed a group effect (BF10 = 3.114; Fig. 1d; Suppl. Table S1), driven by the difference between the CT and TR-K groups (BF10 = 2.116). The win-stay rate similarly showed a group effect (BF10 = 4.226; post-hoc CT vs. TR-K: BF10 = 2.523; Fig. 1e; Suppl. Table S1). However, the loss-shift rate yielded evidence in favor of the null hypothesis regarding a group effect (BF10 = 0.217; Fig. 1f; Suppl. Table S1).

These results suggest that TR-K, but not TR+K, participants exhibit less optimal and more exploratory choices in the task than CT participants.

### Model-Based Measurements

Bayesian Model Selection (BMS) was applied to compare standard Q-learning, Q-learning with decay, and random choice models. The Q-learning with decay model conclusively outperformed the other models across all groups. For the CT group, the Q-learning with decay model achieved an exceedance probability of 1.0000 (Random-Effect analysis (RFX): p(H1|y) = 1.000; Suppl. Fig. S1a, d). The same was true for the TR+K group (exceedance probability = 1.0000; RFX: p(H1|y) = 0.998; Suppl. Fig. S1b, e). In the TR-K group, the Q-learning with decay model dominated with an exceedance probability of 0.9998 (RFX: p(H1|y) = 0.951; Suppl. Fig. S1c, f).

Parameters extracted from the Q-learning with decay model were analyzed via Bayesian ANCOVA. We found strong evidence for a group difference in the estimated learning rate (α) (BF10 = 65.19; Fig. 2a). Post-hoc analyses showed decisive evidence that the learning rate differed between the CT and TR-K groups (BF10 = 649.4), and anecdotal/moderate evidence for a difference between the TR+K and TR-K groups (BF10 = 2.221). Accordingly, the TR-K group exhibited a lower α compared to the other groups, which was distinctly reflected in their learning curves (Fig. 2c-d; Suppl. Fig. S2). In contrast, there was only anecdotal evidence for a group effect on the inverse temperature (β) (BF10 = 1.343; Fig. 2b), and evidence favoring the null hypothesis for the decay rate (γ) (BF10 = 0.190; Fig. 2c).

**Figure 2.**
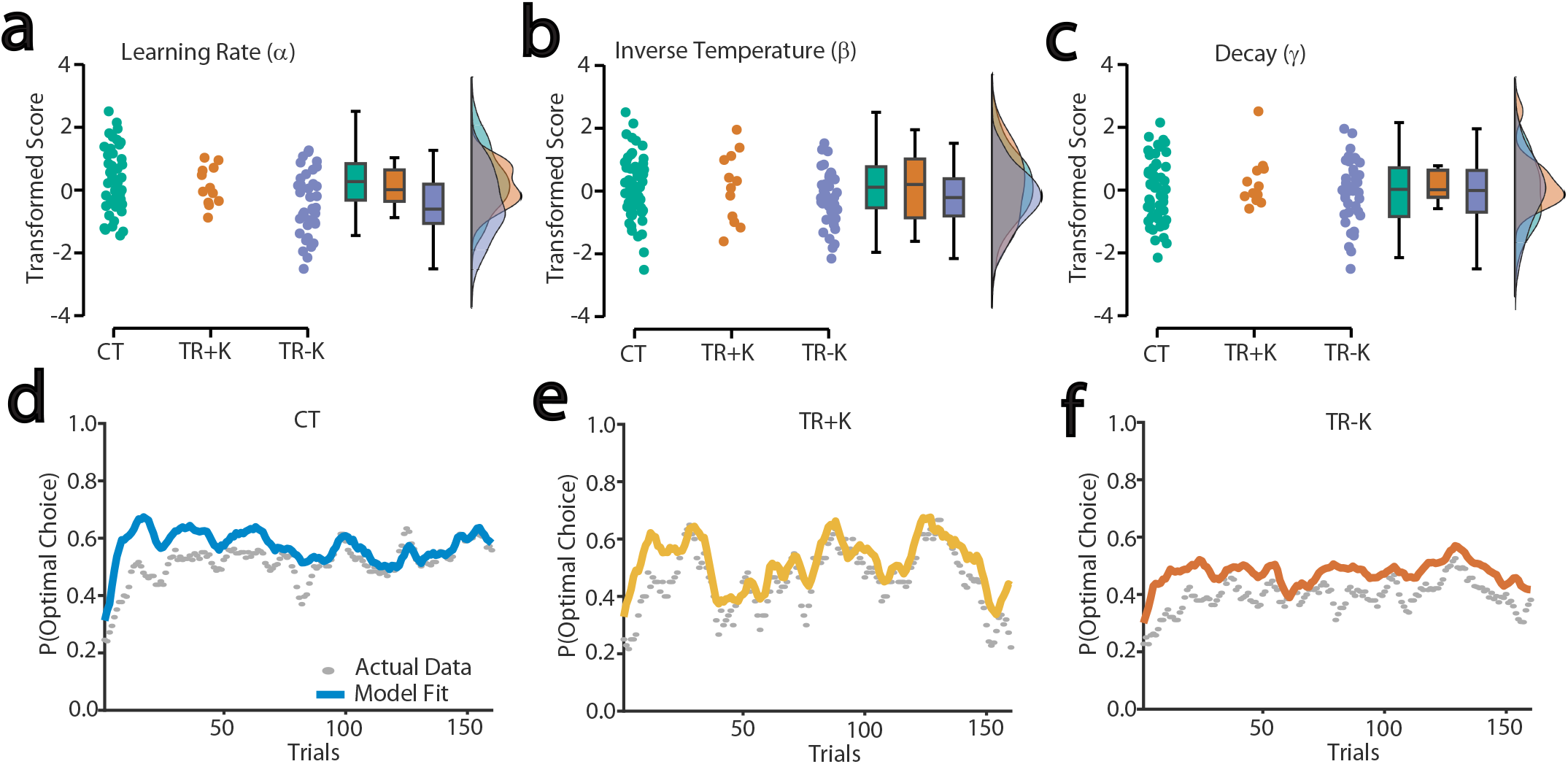
Model-based task measurements in TR+K, TR-K, and CT. a-c, Raincloud plots comparing transformed scores of estimated learning rate α (a), inverse temperature β (b), and decay γ (c) derived from the Q-learning model with decay. **d-f**, Graphs showing the learning curves expressed as changes of optimal choice proportion over 160 trials of the task in CT (d), TR+K (e), and TR-K (f). Solid lines indicate the learning curve with model fitting, and dots indicate actual data, respectively.

The selected model was validated through recovery procedures. Model recovery successfully distinguished the simulated data generation models, yielding posterior probabilities favoring the alternative hypothesis (RFX: p(H1|y) = 1.000 and 0.999 across testing iterations; Suppl. Fig. S3a). Parameter recovery demonstrated high reliability, showing moderate-to-strong cross-correlations between true and estimated parameters on the diagonal (α: r = 0.733; β: r = 0.730; γ: r = 0.785), with weak off-diagonal correlations, indicating minimal parameter collinearity (Suppl. Fig. S3b). Furthermore, simulated behavioral reproduction successfully matched the empirical data, strongly correlating with the observed switch rate (r = 0.741, p = 3.911 × 10^−19^; Suppl. Fig. 3c) and win-stay rate (r = 0.687, p = 1.059 × 10^−15^; Suppl. Fig. 3d).

These results suggest that while all groups utilized a similar RL architecture, TR-K participants exhibited a diminished learning rate and thereby impaired ability to efficiently update expectations based on feedback compared to CT and TR+K participants.

### PFC Responses

To investigate the neural correlates of the RL process, the trial-by-trial Q-value and RPE from the Q-learning model with decay were extracted for each subject in the CT and TR-K groups. These variables were then integrated with the fNIRS data (O2Hb and HHb signals across the 10 measurement locations in the PFC, yielding 20 signal variables) to assess coupling during the task (Suppl. Fig. S4) using the Bayesian LMM.

The Bayesian LMMs demonstrated convergence and stability across all parameters and channels (Suppl. Table S2, S3), with the diagnostic statistic 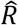 tightly bounded between 1.000 and 1.010 for all fixed and random effects, confirming that the four MCMC chains mixed thoroughly. Furthermore, both bulk and tail Effective Sample Sizes (ESS) were robust (> 500 in all cases, with most exceeding 2,000), indicating high precision in the posterior estimates.

The model revealed credible main effects of RL variables localized to subregions of the PFC. Credible negative estimates for Q-value were observed in the left (R4 O2Hb: Estimate = -0.281, 95% CI [-0.539, -0.025]) and right (R6 HHb: Estimate = -0.185, 95% CI [-0.350, -0.018]) PFrd, and the right PFcdl (R9 HHb: Estimate = -0.242, 95% CI [-0.409, -0.071]; R10 HHb: Estimate = -0.111, 95% CI [-0.220, -0.005]). Assessment of the main effect of Group revealed credible baseline hemodynamic shifts independent of task events (Q-value updates) for the CT group, exhibiting altered baseline O2Hb levels compared to the TR-K group, in the right PFcdm (R7 O2Hb: Estimate = -0.229, 95% CI [-0.441, -0.019]) and PFcdl (R9 O2Hb: Estimate = -0.236, 95% CI [-0.441, -0.032]). In contrast, assessment of the Group modulations of Q-value (Group × Q-value) and RPE (Group × RPE), respectively, revealed that while isolated subregional activity showed marginal variations, the 95% HPDIs for the direct behavioral-hemodynamic interaction contrasts crossed zero across the measured channels.

These results suggest that Q-value and RPE modulate hemodynamic signals in the anterior dorsomedial PFC and right dlPFC; however, such hemodynamic modulations are static metabolic differences, but not the core neural tracking of Q-value and RPE.

## DISCUSSION

In this study, we found that non-kleptomanic theft recidivists demonstrated impaired RL, driven by a diminished learning rate. This underlying neurocomputational deficit caused behavioral inflexibility, evidenced by altered win-stay and switch rates, and resulted in suboptimal decision making characterized by high cumulative regret. In addition, fNIRS evaluation associating these deficits with PFC responses revealed that while trial-by-trial Q-values and RPE modulated hemodynamic signals in specific subregions, such as the anterior dorsomedial PFC and right dorsolateral PFC. Group categorization influenced baseline PFC metabolic states, suggesting an impairment of dismantling the normative value-updating networks utilized during the decision-making task. Collectively, these findings provide a compelling mechanistic explanation for why these non-kleptomanic individuals repeatedly commit thefts despite facing severe, life-altering consequences, such as incarceration.

An intriguing finding in determining the neurobehavioral mechanism of theft recidivism is the significantly lower learning rate identified in the TR-K cohort. In standard RL frameworks, the learning rate dictates how efficiently an individual updates their expectations or Q-values based on new environmental feedback and RPE (Lowet et al., 2020; Schultz et al., 1997; Sutton & Andrew, 2018). A low learning rate indicates that internal expectations are overly rigid. Therefore, these individuals, such as theft recidivists without kleptomania, tend to discount recent feedback and subsequently fail to update their behavior when the environment or payoff contingencies change. This deficit is behaviorally translated into suboptimal decision-making, failing to exploit high-value options, switching behavioral strategies less adaptively, and accumulating significantly more regret over time. In practical, real-world applications, since these individuals possess an inability to update the value of an action based on recent feedback, they fail to build strong internal “values” for stable, safe sources of reward, such as maintaining a job, and avoid behaviors, such as theft, that disrupt that stability. Consequently, they do not properly internalize the long-term rewards of staying out of trouble, leading them to constantly revert to familiar, maladaptive behaviors, i.e., theft. Moreover, they also fail to adequately integrate the severe consequences of punishment of the criminal justice system, such as incarceration, which updates the expected value of stealing to downgrade it to a highly negative outcome and effectively prevent recidivism for individuals with the normal learning rate. However, for theft recidivists without kleptomania, when they return to society, the internal value that they assign to stealing remains unadjusted by their punishment, leading them to reoffend because they literally cannot learn from their mistakes efficiently. Thus, standard punitive measures, which rely entirely on normative value-updating, are therefore inherently ineffective for this specific demographic.

The findings in this study also suggest that recurrent theft does not constitute a singular or neurocognitively homogeneous phenomenon. The TR+K group did not exhibit the same severe deficit in their learning rate as the TR-K group, yielding moderate statistical evidence differentiating the two clinical profiles. Although such insufficient evidence of deficits may be related to a small sample size of the TR+K group, and increasing the sample size may eventually reveal the deficits, individuals with kleptomania appear to possess relatively intact RL systems, such that they likely understand the consequences of their actions and can learn that stealing precipitates negative outcomes. Thus, their recurrent stealing may be tied to other neurobehavioral mechanisms and driven by an overwhelming, condition-specific compulsive urge or a failure of top-down inhibitory control (Blum et al., 2018; Goto et al., 2026; Grant et al., 2007) rather than an inability to learn the fundamental value of the action.

Several major methodological limitations must be considered when interpreting these insights. First, the sample size of the TR+K group was relatively small (n=12), which inherently limits the statistical power of the specific clinical subgroup analysis. Although the application of Bayesian statistical frameworks mitigated this limitation by effectively managing unbalanced data, larger sample sizes would strengthen the group-level inferences. Second, due to strict environmental and logistical constraints within the testing facilities, fNIRS sampling could only be conducted on the CT and TR-K participants. The lack of fNIRS sampling for the TR+K group prevents a comprehensive comparison of PFC hemodynamic variations across the two distinct recidivist subtypes. Finally, the evaluation of populations who had been incarcerated may have inherently introduced a confounding variable related to prolonged institutionalization. Recurrent institutionalization itself could potentially impact the cognitive architecture and cause the observed behavioral and neural alterations.

In conclusion, the current findings contradict the assumptions of the traditional criminal justice system that recurrent theft stems from a moral failing or a calculated, rational choice where the offender actively chooses to ignore the law and that offenders could adjust their future behavior in response to severe structural punishments (Becker, 1968). Recognizing recidivism as, in part, a learning impairment challenges punitive justice models of deterrence and underscores the need for intervention strategies that prioritize targeted cognitive remediation to rely on environmental feedback for rigorous behavioral structuring to bypass their rigid RL systems.

## Acknowledgements

We would like to thank Ms. Miki Kaneda for the assistance, and the staff of the Non-Profit Organization Kurashi O-en Network (Nagoya, Japan), Nishi Hongwanji Byakkoso (Kyoto, Japan), Non-Profit Organization Kyoto MAC (Kyoto, Japan), Kyoto Hogo Ikusei Kai (Kyoto, Japan), and Liberty Women’s House Olive (Otsu, Japan) for recruiting participants, scheduling surveys, and providing various technical supports.

## Data Availability

The data underlying this article will be shared upon reasonable request to the corresponding author.

## Author Contributions/ CRediT Statement

YG contributed to conceptualization, methodology, formal analysis, investigation, data curation, project administration, writing, visualization, validation, supervision, and funding acquisition. SA contributed to conceptualization, resources, and writing. CK contributed to conceptualization, resources, and writing. YAL contributed to formal analysis and writing. All authors have reviewed, edited, and approved the manuscript prior to submission.

## REFERENCES

Adams, R.A., Huys, Q.J., and Roiser, J.P. (2016). Computational Psychiatry: towards a mathematically informed understanding of mental illness. J Neurol Neurosurg Psychiatry 87, 53–63.

Addicott, M.A., Pearson, J.M., Sweitzer, M.M., Barack, D.L., and Platt, M.L. (2017). A Primer on Foraging and the Explore/Exploit Trade-Off for Psychiatry Research. Neuropsychopharmacology 42, 1931–1939.

American Psychiatric Association. (2013). Diagnostic and statistical manual of mental disorders (5th ed.). Washington, D. C.

Asaoka, Y., Won, M., Lee, Y.A., and Goto, Y. (2025). “Neurobehavioral Mechanisms of Kleptomania.,” in Handbook of the Biology and Pathology of Mental Disorders., eds. C.R. Martin, V.R. Preedy, V.B. Patel & R. Rajendram. Springer, Cham.), 10.1007/978-3-031-73368-0_41.

Auzias, G., Coulon, O., and Brovelli, A. (2016). MarsAtlas: A cortical parcellation atlas for functional mapping. Hum Brain Mapp 37, 1573–1592.

Barraclough, D.J., Conroy, M.L., and Lee, D. (2004). Prefrontal cortex and decision making in a mixed-strategy game. Nat Neurosci 7, 404–410.

Becker, G.S. (1968). Crime and punishment: an economic approach. J Polit Econ 76, 169–217.

Blair, R.J.R., Mitchell, D.G.V., Leonard, A., Budhani, S., Peschardt, K.S., and Newman, C. (2004). Passive avoidance learning in individuals with psychopathy: modulation by reward but not by punishment. Pers Individ Differ 37, 1179–1192.

Blanco, C., Grant, J., Petry, N.M., Simpson, H.B., Alegria, A., Liu, S.M., and Hasin, D. (2008). Prevalence and correlates of shoplifting in the United States: results from the National Epidemiologic Survey on Alcohol and Related Conditions (NESARC). Am J Psychiatry 165, 905–913.

Blum, A.W., Odlaug, B.L., Redden, S.A., and Grant, J.E. (2018). Stealing behavior and impulsivity in individuals with kleptomania who have been arrested for shoplifting. Compr Psychiatry 80, 186–191.

Cohen, J.D., Mcclure, S.M., and Yu, A.J. (2007). Should I stay or should I go? How the human brain manages the trade-off between exploitation and exploration. Philos Trans R Soc Lond B Biol Sci 362, 933–942.

Bouneffouf, D., Rish, I., and Cecchi, G.A. (2017). “Bandit models of human behavior: Reward processing in mental disorders.,” in Artificial General Intelligence. AGI 2017. Lecture Notes in Computer Science, eds. E. T., G. B. & A. Potapov. Springer, Cham.).

Daunizeau, J., Adam, V., and Rigoux, L. (2014). VBA: a probabilistic treatment of nonlinear models for neurobiological and behavioural data. PLoS Comput Biol 10, e1003441.

Daw, N.D., O′doherty, J.P., Dayan, P., Seymour, B., and Dolan, R.J. (2006). Cortical substrates for exploratory decisions in humans. Nature 441, 876–879.

Domenech, P., Rheims, S., and Koechlin, E. (2020). Neural mechanisms resolving exploitation-exploration dilemmas in the medial prefrontal cortex. Science 369.

Fazel, S., and Wolf, A. (2015). A Systematic Review of Criminal Recidivism Rates Worldwide: Current Difficulties and Recommendations for Best Practice. PLoS One 10, e0130390.

Goto, Y., Charpentier, R., Yoshino, S., Kita, C., Won, M.J., and Lee, Y.A. (2026). Higher impulsivity in the heterogeneous structure of theft recidivists with and without kleptomania. BioRxiv.

Grant, J.E. (2006). Understanding and treating kleptomania: new models and new treatments. Isr J Psychiatry Relat Sci 43, 81–87.

Grant, J.E., Odlaug, B.L., and Wozniak, J.R. (2007). Neuropsychological functioning in kleptomania. Behav Res Ther 45, 1663–1670.

Harle, K.M., Zhang, S., Schiff, M., Mackey, S., Paulus, M.P., and Yu, A.J. (2015). Altered Statistical Learning and Decision-Making in Methamphetamine Dependence: Evidence from a Two-Armed Bandit Task. Front Psychol 6, 1910.

Hodson, R., Mehta, M., Taylor, S., Lavalley, C.A., Stewart, J.L., Guinjoan, S.M., Ironside, M., White, E.J., Kuplicki, R., Paulus, M.P., and Smith, R. (2026). Attenuated learning rates for negative outcomes in substance use disorders: A replication and extension of prior longitudinal computational modeling results. Drug Alcohol Depend 278, 112922.

Hosking, J.G., Kastman, E.K., Dorfman, H.M., Samanez-Larkin, G.R., Baskin-Sommers, A., Kiehl, K.A., Newman, J.P., and Buckholtz, J.W. (2017). Disrupted Prefrontal Regulation of Striatal Subjective Value Signals in Psychopathy. Neuron 95, 221–231 e224.

JASP Team. (2026). “JASP (Version 0.97.1) [Computer software]”.).

Lee, M.D., and Wagenmakers, E.-J. (2014). Bayesian cognitive modeling: A practical course. Cambridge, UK: Cambridge University Press.

Lowet, A.S., Zheng, Q., Matias, S., Drugowitsch, J., and Uchida, N. (2020). Distributional Reinforcement Learning in the Brain. Trends Neurosci 43, 980–997.

Maia, T.V., and Frank, M.J. (2011). From reinforcement learning models to psychiatric and neurological disorders. Nat Neurosci 14, 154–162.

Montague, P.R., Dolan, R.J., Friston, K.J., and Dayan, P. (2012). Computational psychiatry. Trends Cogn Sci 16, 72–80.

Rouder, J.N., and Morey, R.D. (2012). Default Bayes Factors for Model Selection in Regression. Multivariate Behav Res 47, 877–903.

Rushworth, M.F., Noonan, M.P., Boorman, E.D., Walton, M.E., and Behrens, T.E. (2011). Frontal cortex and reward-guided learning and decision-making. Neuron 70, 1054–1069.

Sazhin, D., Dachs, A., and Smith, D.V. (2025). Meta-Analysis Reveals That Explore-Exploit Decisions are Dissociable by Activation in the Dorsal Lateral Prefrontal Cortex, Anterior Insula, and the Dorsal Anterior Cingulate Cortex. bioRxiv.

Schultz, W., Dayan, P., and Montague, P.R. (1997). A neural substrate of prediction and reward. Science 275, 1593–1599.

Seo, H., Barraclough, D.J., and Lee, D. (2007). Dynamic signals related to choices and outcomes in the dorsolateral prefrontal cortex. Cereb Cortex 17 Suppl 1, i110–117.

Seymour, B., Daw, N.D., Roiser, J.P., Dayan, P., and Dolan, R. (2012). Serotonin selectively modulates reward value in human decision-making. J Neurosci 32, 5833–5842.

Speekenbrink, M., and Konstantinidis, E. (2015). Uncertainty and exploration in a restless bandit problem. Top Cogn Sci 7, 351–367.

Sutton, R.S., and Barto, A.G. (2018). Reinforcement learning: An introduction. Cambridge, MA: MIT Press.

Wiehler, A., Chakroun, K., and Peters, J. (2021). Attenuated Directed Exploration during Reinforcement Learning in Gambling Disorder. J Neurosci 41, 2512–2522.

Yaniv, G. (2009). Shoplifting, monitoring and price determination. J Socio-Econ 38, 608–610.

Yukhnenko, D., Farouki, L., and Fazel, S. (2023). Criminal recidivism rates globally: A 6-year systematic review update. J Crim Justice 88, 102115.

