## Supplementary Materials for "Impaired Reinforcement Learning Underlying Explore-Exploit Decision Making in Theft Recidivists"

**Supplementary Methods**

**Supplementary Table S1**
A summary of the model-free and model-based measurements with 4-arm Bandit Task.

**Supplementary Table S2**
A summary of the Bayesian LMM results for associations of Q-values with PFC responses.

**Supplementary Table S3**
A summary of the Bayesian LMM results for associations of RPE with PFC responses.

**Supplementary Figure S1**
Group-level Bayesian Model Selection (BMS).

**Supplementary Figure S2**
An example of the model evaluation of a single participant's choices.

**Supplementary Figure S3**
Validations of the model with model and parameter recovery analyses and behavioral reproduction.

**Supplementary Figure S4**
A single subject example of fNIRS signals with model-derived latent reinforcement variables.

### SUPPLEMENTARY METHODS

#### Model-free Behavioral Metrics in 4-arm Bandit Task

Model-free behavioral metrics, such as cumulative regret, optimal choice proportion, exploitative choice proportion, switch rate, and win-stay/lose-shift rates, were calculated with the following mathematical definitions;

*(Empirical) Cumulative regret:*

$$CR = \sum_{i=1}^{T} (\mu^* - R_i)$$

*Optimal choice proportion:*

$$P_{opt} = \frac{1}{T}\sum_{i=1}^{T} \mathbb{I}\,(C_i = C^*)$$

*Exploitative choice proportion:*

$$P_{exploit} = \frac{1}{T}\sum_{i=1}^{T} \mathbb{I}\left(C_i = \arg\max_{c\in C} \hat{\mu}_i(c)\right)$$

*Switch rate:*

$$SR = \frac{1}{T-1}\sum_{i=2}^{T} \mathbb{I}\,(C_i \neq C_{i-1})$$

*(Chance-corrected) win-stay rate:*

$$WS = \frac{\sum_{i=1}^{T-1} \mathbb{I}\,(C_{i+1} = C_i) \cdot \mathbb{I}(R_i = 1)}{\sum_{i=1}^{T-1} \mathbb{I}\,(R_i = 1)} - 0.25$$

*(Chance-corrected)* lose-shift rate:

$$LS = \frac{\sum_{i=1}^{T-1} \mathbb{I}\,(C_{i+1} \neq C_i) \cdot \mathbb{I}(R_i = 0)}{\sum_{i=1}^{T-1} \mathbb{I}\,(R_i = 0)} - 0.75$$

, where $C_i$ is the choice made on trial i, $R_i$ is the actual reward received on trial i, C* is the true optimal choice, $\mu(C_i)$ is the true expected reward of the choice made on trial i, $\mu^*$ is the true expected reward of the optimal choice C*, and $\hat{\mu}_i(c)$ is the participant's internal estimated value for a given choice c at the exact moment before making a decision on trial i. In win-stay/lose shift rate, $R_i \in \{0,1\}$, where 1 is a win (higher than the previous payoff) and 0 is a lose (lower than the previous payoff).

#### Data Analysis for Model-based Measurements

*Computational Modeling:* To evaluate the latent cognitive mechanisms driving choice behavior, three distinct models were fitted to the trial-by-trial sequence of choices and rewards for each participant.

*(Model 1: Standard Q-Learning Model)* This model assumes that participants learn the expected value (Q-value) of each arm through a standard Rescorla-Wagner prediction error update. The Q-value for the chosen arm c at trial t is updated as follows:

$$Q_{t+1}(c) = Q_t(c) + \alpha\big(R_t - Q_t(c)\big)$$

, where $\alpha \in (0,1)$ is the learning rate governing the weight of the prediction error, and $R_t$ is the reward received. The Q-values of the unchosen arms remain unchanged.

*(Model 2: Q-Learning with Decay Model)* This model introduces an unchosen-arm decay mechanism, assuming that the valuation of unexplored options regresses toward a

neutral prior. The chosen arm is updated identically to that in Model 1. However, the Q-values for the unchosen arms u decay toward 0.5 (the midpoint of the normalized reward space):

$$Q_{t+1}(u) = (1 - \delta)Q_t(u) + \delta(0.5)$$

, where δ ∈ (0,1) is the decay rate (analyzed as γ in group comparisons).

*(Model 3: Random Choice Model)* A random choice model was included as a baseline to evaluate whether participants' behavior was better explained by reinforcement learning or random guessing. This model assumes that choices are made independently of the past reward history. No internal state or Q-values are updated, and the probability of selecting any of the four arms remains fixed at the chance level for every trial:

$$\mathrm{P}(\mathrm{a_t}) = 0.25$$

*(Action Selection: Softmax)* For Models 1 and 2, choice probabilities were generated using a softmax action selection rule:

$$P(c_t = i) = \frac{\exp(\beta Q_t(i)))}{\sum_{j=1}^{4} \exp(\beta Q_t(j)))}$$

, where β is the inverse temperature parameter regulating exploration versus exploitation. To ensure numerical stability, the input Q-values were mean-centered to the maximum current Q-value prior to exponentiation.

*Parameter Estimation:* Parameters were estimated for each participant using Variational Bayes within the nonlinear state-space framework of the VBA toolbox. To enforce theoretically valid parameter ranges without relying on hard boundary constraints during optimization, native parameters were estimated in unbounded space and mapped via logistic sigmoid transformations:

*(Learning Rate α and Decay γ)* Transformed via standard sigmoid bounds to restrict values to (0,1):

$$\theta = \frac{1}{1 + \exp(-P)}$$

*(Inverse Temperature β)* Bounded between 0 and 20 to prevent computational explosion:

$$\beta = \frac{20}{1 + \exp(-P)}$$

For model estimation, prior parameter means were set to 0 with a variance of 3, except for the β prior, which was shifted to yield an initial expectation space optimized for the observed reward variance.

*Model Validation Framework:* A comprehensive validation pipeline was executed to verify the robustness and interpretability of the winning model.

*(Model Recovery Analysis)* To ensure that the BMS procedure could reliably distinguish between standard Q-learning (Model 1) and Q-learning with decay (Model 2), a cross-fitting model recovery analysis was conducted. 25 synthetic datasets were generated per model using a drifting random walk environment ($\lambda = 0.98$, $\theta = 50$, $\sigma_d = 2.8$, $\sigma_o = 4$). Both models were then refitted to both sets of synthetic data. A confusion matrix of the resulting Exceedance Probabilities confirmed the framework's capacity to accurately identify the true generative model without false-positive overfitting.

*(Parameter Recovery Analysis)* To confirm that the winning model's parameters (α, β, γ) were independently identifiable, 100 synthetic datasets were generated using known parameter values sampled from the priors. The synthetic data were fitted using the VBA procedure, and Pearson correlation coefficients were calculated between the true generative parameters and the estimated parameters. Furthermore, parameter cross-correlations (e.g., true α vs. estimated β) were evaluated via a correlation heatmap to rule out problematic collinearity or trade-offs between the learning rate, decay, and exploration parameters.

**Supplementary Table S1. A summary of the model-free and model-based measurements with 4-arm Bandit Task.**

| | | CT (n=53) | TR+K (n=12) | TR-K (n=38) |
|---|---|---|---|---|
| **Model-free** | **Cumulative Regret** | 9.523 ± 0.676 | 11.467± 1.928 | 14.288 ± 0.915 |
| | **Optimal Choice** | 0.513 ± 0.022 | 0.462 ± 0.039 | 0.391 ± 0.027 |
| | **Exploitative Choice** | 0.609 ± 0.026 | 0.536 ± 0.053 | 0.450 ± 0.033 |
| | **Switch Rate** | 0.494 ± 0.035 | 0.513 ± 0.060 | 0.644 ± 0.048 |
| | **Win-stay Rate** | 0.333 ± 0.038 | 0.322 ± 0.070 | 0.158 ± 0.053 |
| | **Lose-shift Rate** | 0.113 ± 0.023 | 0.070 ± 0.054 | 0.049 ± 0.041 |
| **Model-based** | **Learning Rate (α)** | 0.481 ± 0.036 | 0.414 ± 0.059 | 0.267 ± 0.037 |
| | **Inverse Temperature (β)** | 12.085 ± 0.715 | 12.074 ± 1.649 | 10.212 ± 0.785 |
| | **Decay (γ)** | 0.395 ± 0.038 | 0.449 ± 0.056 | 0.390 ± 0.042 |
| | **McFadden's pseudo-$R^2$** | 0.298 ± 0.029 | 0.245 ± 0.045 | 0.166 ± 0.034 |
| | **BIC (Decay Model)** | 325.6 ± 12.63 | 340.1 ± 20.48 | 380.5 ± 14.77 |
| | **BIC (Standard Model)** | 334.4 ± 12.15 | 359.3 ± 17.01 | 375.9 ± 16.36 |
| | **BIC (Random Model)** | 442.2 ± 0.339 | 430.0 ± 5.192 | 438.2 ± 1.817 |

CT – Control subjects with no criminal record; TR+K – Theft recidivist with kleptomania; TR-K –Theft recidivist without kleptomania; BIC – Bayesian information criteria; Model-based metrics (except BIC with standard Q-learning model and random choice model) are derived with the Q-learning with decay model. The data are represented as mean ± s.e.m.

**Supplementary Table S2. A summary of the Bayesian LMM results for associations of Q-values with PFC responses.**

**Estimates (difference from Intercept for Group, Q-value, and Group x Q-value)**

| Site | Signal | Term | Level | Estimate (β) | s.e.m. | 95% CI | | $\hat{R}$ | ESS (bulk) | ESS (tail) |
|---|---|---|---|---|---|---|---|---|---|---|
| | | | | | | Lower | Upper | | | |
| R1 | O2Hb | Intercept | | 0.048 | 0.083 | -0.118 | 0.219 | 1.006 | 546.2 | 1268 |
| R1 | O2Hb | Group | TR-K | 0.108 | 0.089 | -0.073 | 0.290 | 1.005 | 727.1 | 1343 |
| R1 | O2Hb | Group | CT | -0.137 | 0.084 | -0.311 | 0.034 | 1.004 | 717.0 | 1194 |
| R1 | O2Hb | Q-value | | -0.046 | 0.046 | -0.137 | 0.046 | 1.003 | 1533 | 4582 |
| R1 | O2Hb | Group x Q-value | TR-K | 0.004 | 0.074 | -0.144 | 0.152 | 1.001 | 1901 | 5287 |
| R1 | O2Hb | Group x Q-value | CT | -0.096 | 0.053 | -0.201 | 0.010 | 1.003 | 1539 | 4486 |
| R1 | HHb | Intercept | | 0.045 | 0.080 | -0.119 | 0.196 | 1.004 | 632.1 | 1164 |
| R1 | HHb | Group | TR-K | 0.055 | 0.088 | -0.112 | 0.222 | 1.003 | 757.2 | 2149 |
| R1 | HHb | Group | CT | -0.100 | 0.085 | -0.263 | 0.060 | 1.003 | 691.4 | 1777 |
| R1 | HHb | Q-value | | -0.071 | 0.056 | -0.181 | 0.039 | 1.000 | 3704 | 8339 |
| R1 | HHb | Group x Q-value | TR-K | -0.067 | 0.092 | -0.245 | 0.118 | 1.000 | 4314 | 7562 |
| R1 | HHb | Group x Q-value | CT | -0.076 | 0.064 | -0.200 | 0.049 | 1.001 | 3125 | 6845 |
| R2 | O2Hb | Intercept | | 0.035 | 0.084 | -0.127 | 0.202 | 1.002 | 813.9 | 1798 |
| R2 | O2Hb | Group | TR-K | 0.081 | 0.080 | -0.077 | 0.242 | 1.002 | 1314 | 2394 |
| R2 | O2Hb | Group | CT | -0.148 | 0.079 | -0.304 | 0.009 | 1.003 | 1240 | 2247 |
| R2 | O2Hb | Q-value | | -0.105 | 0.107 | -0.317 | 0.100 | 1.002 | 1966 | 4361 |
| R2 | O2Hb | Group x Q-value | TR-K | -0.191 | 0.171 | -0.528 | 0.148 | 1.001 | 2242 | 4910 |
| R2 | O2Hb | Group x Q-value | CT | -0.020 | 0.128 | -0.274 | 0.237 | 1.003 | 1613 | 3243 |
| R2 | HHb | Intercept | | 0.039 | 0.087 | -0.133 | 0.209 | 1.003 | 1120 | 2354 |
| R2 | HHb | Group | TR-K | -0.068 | 0.084 | -0.235 | 0.099 | 1.007 | 1302 | 2736 |
| R2 | HHb | Group | CT | -0.055 | 0.084 | -0.221 | 0.112 | 1.008 | 1323 | 2453 |
| R2 | HHb | Q-value | | -0.193 | 0.101 | -0.396 | 0.010 | 1.001 | 2486 | 4738 |
| R2 | HHb | Group x Q-value | TR-K | -0.355 | 0.160 | -0.675 | -0.034 | 1.002 | 2923 | 5673 |
| R2 | HHb | Group x Q-value | CT | -0.032 | 0.119 | -0.273 | 0.203 | 1.001 | 1900 | 4043 |
| R3 | O2Hb | Intercept | | 0.049 | 0.074 | -0.099 | 0.198 | 1.004 | 1236 | 2162 |
| R3 | O2Hb | Group | TR-K | 0.042 | 0.089 | -0.137 | 0.217 | 1.003 | 1376 | 3064 |
| R3 | O2Hb | Group | CT | -0.118 | 0.082 | -0.278 | 0.045 | 1.004 | 1247 | 2610 |
| R3 | O2Hb | Q-value | | -0.120 | 0.090 | -0.295 | 0.061 | 1.001 | 3990 | 6542 |
| R3 | O2Hb | Group x Q-value | TR-K | -0.223 | 0.147 | -0.506 | 0.070 | 1.001 | 4524 | 7402 |
| R3 | O2Hb | Group x Q-value | CT | -0.016 | 0.104 | -0.222 | 0.191 | 1.002 | 3018 | 5716 |
| R3 | HHb | Intercept | | 0.020 | 0.077 | -0.127 | 0.168 | 1.004 | 830.6 | 2017 |
| R3 | HHb | Group | TR-K | 0.079 | 0.095 | -0.112 | 0.263 | 1.004 | 1290 | 2514 |
| R3 | HHb | Group | CT | -0.080 | 0.086 | -0.253 | 0.092 | 1.004 | 1007 | 1866 |
| R3 | HHb | Q-value | | -0.002 | 0.119 | -0.235 | 0.233 | 1.001 | 2110 | 4421 |
| R3 | HHb | Group x Q-value | TR-K | 0.042 | 0.190 | -0.328 | 0.413 | 1.001 | 2683 | 5296 |
| R3 | HHb | Group x Q-value | CT | -0.046 | 0.145 | -0.332 | 0.244 | 1.002 | 1455 | 2820 |
| R4 | O2Hb | Intercept | | 0.063 | 0.080 | -0.093 | 0.223 | 1.001 | 836.1 | 2174 |
| R4 | O2Hb | Group | TR-K | -0.046 | 0.100 | -0.242 | 0.159 | 1.004 | 1493 | 2618 |
| R4 | O2Hb | Group | CT | -0.133 | 0.092 | -0.316 | 0.046 | 1.002 | 1172 | 2465 |
| R4 | O2Hb | Q-value | | -0.281 | 0.132 | -0.539 | -0.025 | 1.001 | 2146 | 4343 |
| R4 | O2Hb | Group x Q-value | TR-K | -0.471 | 0.214 | -0.891 | -0.058 | 1.001 | 2773 | 5151 |
| R4 | O2Hb | Group x Q-value | CT | -0.091 | 0.153 | -0.407 | 0.219 | 1.001 | 1509 | 3050 |
| R4 | HHb | Intercept | | 0.084 | 0.068 | -0.052 | 0.215 | 1.009 | 623.6 | 1627 |
| R4 | HHb | Group | TR-K | -0.215 | 0.130 | -0.467 | 0.047 | 1.004 | 1044 | 1897 |
| R4 | HHb | Group | CT | -0.076 | 0.106 | -0.298 | 0.133 | 1.005 | 545.7 | 1165 |
| R4 | HHb | Q-value | | -0.458 | 0.241 | -0.950 | 0.029 | 1.007 | 759.8 | 1488 |
| R4 | HHb | Group x Q-value | TR-K | -0.853 | 0.398 | -1.635 | -0.091 | 1.006 | 1071 | 2091 |
| R4 | HHb | Group x Q-value | CT | -0.063 | 0.301 | -0.67 | 0.514 | 1.005 | 579.8 | 1305 |
| R5 | O2Hb | Intercept | | -0.008 | 0.083 | -0.174 | 0.160 | 1.002 | 870.2 | 1944 |
| R5 | O2Hb | Group | TR-K | -0.034 | 0.086 | -0.204 | 0.138 | 1.001 | 1118 | 2193 |
| R5 | O2Hb | Group | CT | 0.032 | 0.084 | -0.132 | 0.199 | 1.003 | 1008 | 1845 |
| R5 | O2Hb | Q-value | | -0.003 | 0.115 | -0.230 | 0.221 | 1.003 | 1564 | 2916 |
| R5 | O2Hb | Group x Q-value | TR-K | -0.157 | 0.181 | -0.526 | 0.207 | 1.003 | 1807 | 3961 |
| R5 | O2Hb | Group x Q-value | CT | 0.150 | 0.135 | -0.118 | 0.415 | 1.003 | 1241 | 2585 |
| R5 | HHb | Intercept | | 0.033 | 0.077 | -0.121 | 0.183 | 1.002 | 1227 | 2679 |
| R5 | HHb | Group | TR-K | 0.025 | 0.087 | -0.149 | 0.200 | 1.004 | 1828 | 3585 |

| | | | | | | | | | | |
|---|---|---|---|---|---|---|---|---|---|---|
| R5 | HHb | Group | CT | -0.071 | 0.082 | -0.229 | 0.092 | 1.003 | 1514 | 2623 |
| R5 | HHb | Q-value | | -0.072 | 0.100 | -0.266 | 0.126 | 1.000 | 4269 | 7622 |
| R5 | HHb | Group x Q-value | TR-K | -0.181 | 0.166 | -0.504 | 0.136 | 1.001 | 4949 | 8173 |
| R5 | HHb | Group x Q-value | CT | 0.037 | 0.115 | -0.191 | 0.267 | 1.001 | 3312 | 5476 |
| R6 | O2Hb | Intercept | | 0.038 | 0.084 | -0.131 | 0.207 | 1.001 | 691.4 | 1548 |
| R6 | O2Hb | Group | TR-K | -0.066 | 0.095 | -0.254 | 0.117 | 1.007 | 954.0 | 2239 |
| R6 | O2Hb | Group | CT | -0.025 | 0.088 | -0.199 | 0.154 | 1.007 | 855.9 | 1781 |
| R6 | O2Hb | Q-value | | -0.144 | 0.094 | -0.330 | 0.041 | 1.001 | 2456 | 4450 |
| R6 | O2Hb | Group x Q-value | TR-K | -0.126 | 0.150 | -0.428 | 0.175 | 1.001 | 2963 | 5227 |
| R6 | O2Hb | Group x Q-value | CT | -0.163 | 0.110 | -0.387 | 0.053 | 1.002 | 1753 | 3269 |
| R6 | HHb | Intercept | | 0.031 | 0.087 | -0.143 | 0.211 | 1.003 | 766.4 | 1757 |
| R6 | HHb | Group | TR-K | -0.126 | 0.095 | -0.314 | 0.059 | 1.004 | 972.1 | 1794 |
| R6 | HHb | Group | CT | 0.008 | 0.090 | -0.168 | 0.191 | 1.005 | 859.0 | 1522 |
| R6 | HHb | Q-value | | -0.185 | 0.085 | -0.350 | -0.018 | 1.001 | 3019 | 5952 |
| R6 | HHb | Group x Q-value | TR-K | -0.212 | 0.137 | -0.481 | 0.065 | 1.001 | 3667 | 6636 |
| R6 | HHb | Group x Q-value | CT | -0.158 | 0.099 | -0.350 | 0.032 | 1.001 | 2332 | 4965 |
| R7 | O2Hb | Intercept | | 0.065 | 0.074 | -0.08 | 0.212 | 1.005 | 1541 | 2809 |
| R7 | O2Hb | Group | TR-K | 0.124 | 0.084 | -0.048 | 0.295 | 1.003 | 1757 | 3220 |
| R7 | O2Hb | Group | CT | -0.169 | 0.080 | -0.329 | -0.011 | 1.002 | 1653 | 2986 |
| R7 | O2Hb | Q-value | | -0.070 | 0.077 | -0.221 | 0.084 | 1.000 | 5851 | 8262 |
| R7 | O2Hb | Group x Q-value | TR-K | -0.119 | 0.127 | -0.368 | 0.130 | 1.000 | 6395 | 8230 |
| R7 | O2Hb | Group x Q-value | CT | -0.021 | 0.088 | -0.197 | 0.155 | 1.000 | 5086 | 8052 |
| R7 | HHb | Intercept | | 0.014 | 0.077 | -0.140 | 0.165 | 1.003 | 1342 | 3121 |
| R7 | HHb | Group | TR-K | -0.086 | 0.086 | -0.258 | 0.081 | 1.001 | 1851 | 3327 |
| R7 | HHb | Group | CT | 0.042 | 0.081 | -0.115 | 0.205 | 1.002 | 1714 | 2959 |
| R7 | HHb | Q-value | | -0.070 | 0.078 | -0.222 | 0.089 | 1.001 | 4911 | 8004 |
| R7 | HHb | Group x Q-value | TR-K | -0.075 | 0.127 | -0.325 | 0.186 | 1.000 | 5144 | 7511 |
| R7 | HHb | Group x Q-value | CT | -0.065 | 0.090 | -0.243 | 0.114 | 1.001 | 4517 | 7600 |
| R8 | O2Hb | Intercept | | -0.017 | 0.084 | -0.184 | 0.154 | 1.004 | 675.4 | 1334 |
| R8 | O2Hb | Group | TR-K | -0.093 | 0.102 | -0.294 | 0.107 | 1.003 | 848.6 | 1872 |
| R8 | O2Hb | Group | CT | 0.063 | 0.092 | -0.118 | 0.251 | 1.006 | 769.5 | 1525 |
| R8 | O2Hb | Q-value | | -0.047 | 0.129 | -0.305 | 0.207 | 1.002 | 1359 | 2505 |
| R8 | O2Hb | Group x Q-value | TR-K | 0.045 | 0.210 | -0.371 | 0.453 | 1.003 | 1725 | 3475 |
| R8 | O2Hb | Group x Q-value | CT | -0.139 | 0.156 | -0.451 | 0.173 | 1.009 | 1040 | 2149 |
| R8 | HHb | Intercept | | 0.025 | 0.082 | -0.133 | 0.189 | 1.003 | 922.5 | 1754 |
| R8 | HHb | Group | TR-K | -0.147 | 0.113 | -0.377 | 0.081 | 1.004 | 1499 | 2695 |
| R8 | HHb | Group | CT | 0.021 | 0.103 | -0.182 | 0.228 | 1.004 | 874.1 | 1732 |
| R8 | HHb | Q-value | | -0.197 | 0.185 | -0.561 | 0.156 | 1.002 | 1458 | 2977 |
| R8 | HHb | Group x Q-value | TR-K | -0.281 | 0.296 | -0.862 | 0.301 | 1.003 | 1784 | 3123 |
| R8 | HHb | Group x Q-value | CT | -0.113 | 0.220 | -0.552 | 0.327 | 1.003 | 853.7 | 1854 |
| R9 | O2Hb | Intercept | | 0.128 | 0.078 | -0.022 | 0.275 | 1.007 | 746.2 | 1761 |
| R9 | O2Hb | Group | TR-K | 0.014 | 0.105 | -0.198 | 0.233 | 1.003 | 1100 | 2046 |
| R9 | O2Hb | Group | CT | -0.203 | 0.098 | -0.392 | -0.011 | 1.004 | 794.1 | 2040 |
| R9 | O2Hb | Q-value | | -0.297 | 0.185 | -0.646 | 0.074 | 1.005 | 964.0 | 2288 |
| R9 | O2Hb | Group x Q-value | TR-K | -0.481 | 0.294 | -1.055 | 0.116 | 1.001 | 1315 | 2652 |
| R9 | O2Hb | Group x Q-value | CT | -0.112 | 0.227 | -0.554 | 0.333 | 1.001 | 749.3 | 1643 |
| R9 | HHb | Intercept | | 0.069 | 0.079 | -0.090 | 0.228 | 1.004 | 1446 | 2655 |
| R9 | HHb | Group | TR-K | -0.088 | 0.082 | -0.254 | 0.075 | 1.001 | 2035 | 3842 |
| R9 | HHb | Group | CT | -0.066 | 0.080 | -0.221 | 0.098 | 1.001 | 1735 | 3062 |
| R9 | HHb | Q-value | | -0.242 | 0.084 | -0.409 | -0.071 | 1.001 | 4939 | 8330 |
| R9 | HHb | Group x Q-value | TR-K | -0.377 | 0.138 | -0.647 | -0.101 | 1.000 | 5235 | 8985 |
| R9 | HHb | Group x Q-value | CT | -0.107 | 0.099 | -0.306 | 0.089 | 1.002 | 3946 | 6610 |
| R10 | O2Hb | Intercept | | 0.021 | 0.064 | -0.105 | 0.149 | 1.002 | 1140 | 1910 |
| R10 | O2Hb | Group | TR-K | 0.044 | 0.082 | -0.118 | 0.212 | 1.002 | 1215 | 2063 |
| R10 | O2Hb | Group | CT | -0.039 | 0.073 | -0.190 | 0.110 | 1.003 | 1139 | 2240 |
| R10 | O2Hb | Q-value | | 0.007 | 0.070 | -0.133 | 0.147 | 1.001 | 3591 | 7162 |
| R10 | O2Hb | Group x Q-value | TR-K | 0.107 | 0.116 | -0.117 | 0.339 | 1.001 | 3560 | 7050 |
| R10 | O2Hb | Group x Q-value | CT | -0.093 | 0.080 | -0.251 | 0.064 | 1.001 | 3143 | 7350 |
| R10 | HHb | Intercept | | 0.031 | 0.058 | -0.087 | 0.143 | 1.003 | 1361 | 2561 |
| R10 | HHb | Group | TR-K | -0.075 | 0.077 | -0.227 | 0.077 | 1.002 | 1729 | 3216 |
| R10 | HHb | Group | CT | 0.004 | 0.071 | -0.138 | 0.146 | 1.001 | 1510 | 3005 |
| R10 | HHb | Q-value | | -0.111 | 0.054 | -0.220 | -0.005 | 1.002 | 2186 | 3906 |
| R10 | HHb | Group x Q-value | TR-K | -0.168 | 0.088 | -0.341 | 0.004 | 1.001 | 3102 | 6386 |

| | | | | | | | | | | |
|---|---|---|---|---|---|---|---|---|---|---|
| R10 | HHb | Group x Q-value | CT | -0.055 | 0.064 | -0.185 | 0.071 | 1.001 | 1765 | 3322 |

**Fixed Effects Estimates**

| Site | Signal | Term | Estimate (β) | s.e.m. | 95% CI | | $\hat{R}$ | ESS |
|---|---|---|---|---|---|---|---|---|
| | | | | | Lower | Upper | | |
| R1 | O2Hb | Intercept | 0.048 | 0.085 | -0.118 | 0.219 | 1.006 | 532.9 |
| R1 | O2Hb | Group | -0.151 | 0.114 | -0.377 | 0.073 | 1.005 | 703.9 |
| R1 | O2Hb | Q-value | -0.046 | 0.046 | -0.137 | 0.046 | 1.003 | 1482 |
| R1 | O2Hb | Group x Q-value | -0.071 | 0.065 | -0.198 | 0.058 | 1.001 | 1960 |
| R1 | HHb | Intercept | 0.045 | 0.080 | -0.119 | 0.196 | 1.004 | 635.5 |
| R1 | HHb | Group | -0.107 | 0.109 | -0.320 | 0.102 | 1.004 | 704.5 |
| R1 | HHb | Q-value | -0.071 | 0.056 | -0.181 | 0.039 | 1.000 | 3678 |
| R1 | HHb | Group x Q-value | -0.006 | 0.080 | -0.167 | 0.149 | 1.000 | 3933 |
| R2 | O2Hb | Intercept | 0.035 | 0.084 | -0.127 | 0.202 | 1.002 | 811.4 |
| R2 | O2Hb | Group | -0.200 | 0.118 | -0.439 | 0.029 | 1.003 | 1100 |
| R2 | O2Hb | Q-value | -0.105 | 0.107 | -0.317 | 0.100 | 1.002 | 1958 |
| R2 | O2Hb | Group x Q-value | 0.121 | 0.152 | -0.169 | 0.426 | 1.002 | 1999 |
| R2 | HHb | Intercept | 0.039 | 0.087 | -0.133 | 0.209 | 1.003 | 1112 |
| R2 | HHb | Group | -0.063 | 0.125 | -0.303 | 0.186 | 1.001 | 1032 |
| R2 | HHb | Q-value | -0.193 | 0.103 | -0.396 | 0.010 | 1.000 | 2482 |
| R2 | HHb | Group x Q-value | 0.229 | 0.142 | -0.054 | 0.509 | 1.003 | 2319 |
| R3 | O2Hb | Intercept | 0.049 | 0.075 | -0.099 | 0.198 | 1.004 | 1230 |
| R3 | O2Hb | Group | -0.159 | 0.104 | -0.362 | 0.047 | 1.003 | 1098 |
| R3 | O2Hb | Q-value | -0.120 | 0.091 | -0.295 | 0.061 | 1.001 | 3984 |
| R3 | O2Hb | Group x Q-value | 0.146 | 0.127 | -0.109 | 0.391 | 1.001 | 3791 |
| R3 | HHb | Intercept | 0.020 | 0.076 | -0.127 | 0.168 | 1.004 | 825.0 |
| R3 | HHb | Group | -0.093 | 0.108 | -0.303 | 0.122 | 1.006 | 1018 |
| R3 | HHb | Q-value | -0.002 | 0.120 | -0.235 | 0.233 | 1.001 | 2104 |
| R3 | HHb | Group x Q-value | -0.062 | 0.170 | -0.391 | 0.269 | 1.000 | 1953 |
| R4 | O2Hb | Intercept | 0.063 | 0.080 | -0.093 | 0.223 | 1.001 | 833.4 |
| R4 | O2Hb | Group | -0.147 | 0.116 | -0.378 | 0.081 | 1.004 | 1093 |
| R4 | O2Hb | Q-value | -0.281 | 0.132 | -0.539 | -0.025 | 1.001 | 2141 |
| R4 | O2Hb | Group x Q-value | 0.269 | 0.188 | -0.096 | 0.645 | 1.000 | 2132 |
| R4 | HHb | Intercept | 0.084 | 0.068 | -0.052 | 0.215 | 1.009 | 611.0 |
| R4 | HHb | Group | -0.079 | 0.096 | -0.267 | 0.103 | 1.006 | 706.4 |
| R4 | HHb | Q-value | -0.458 | 0.249 | -0.950 | 0.029 | 1.007 | 724.1 |
| R4 | HHb | Group x Q-value | 0.558 | 0.355 | -0.130 | 1.252 | 1.004 | 856.1 |
| R5 | O2Hb | Intercept | -0.008 | 0.085 | -0.174 | 0.160 | 1.002 | 866.0 |
| R5 | O2Hb | Group | -0.023 | 0.121 | -0.258 | 0.214 | 1.003 | 823.2 |
| R5 | O2Hb | Q-value | -0.003 | 0.115 | -0.230 | 0.221 | 1.003 | 1561 |
| R5 | O2Hb | Group x Q-value | 0.217 | 0.162 | -0.101 | 0.534 | 1.003 | 1576 |
| R5 | HHb | Intercept | 0.033 | 0.078 | -0.121 | 0.183 | 1.002 | 1216 |
| R5 | HHb | Group | -0.117 | 0.107 | -0.326 | 0.097 | 1.005 | 1316 |
| R5 | HHb | Q-value | -0.072 | 0.100 | -0.266 | 0.126 | 1.000 | 4260 |
| R5 | HHb | Group x Q-value | 0.154 | 0.141 | -0.121 | 0.434 | 1.000 | 4220 |
| R6 | O2Hb | Intercept | 0.038 | 0.086 | -0.131 | 0.207 | 1.001 | 700.1 |
| R6 | O2Hb | Group | 0.037 | 0.117 | -0.187 | 0.269 | 1.009 | 796.1 |
| R6 | O2Hb | Q-value | -0.144 | 0.095 | -0.330 | 0.041 | 1.001 | 2440 |
| R6 | O2Hb | Group x Q-value | -0.026 | 0.133 | -0.285 | 0.236 | 1.001 | 2395 |
| R6 | HHb | Intercept | 0.031 | 0.089 | -0.143 | 0.211 | 1.003 | 760.3 |
| R6 | HHb | Group | 0.083 | 0.126 | -0.160 | 0.338 | 1.006 | 787.5 |
| R6 | HHb | Q-value | -0.185 | 0.085 | -0.350 | -0.018 | 1.001 | 3001 |
| R6 | HHb | Group x Q-value | 0.038 | 0.120 | -0.199 | 0.270 | 1.000 | 3305 |
| R7 | O2Hb | Intercept | 0.065 | 0.075 | -0.080 | 0.212 | 1.005 | 1554 |
| R7 | O2Hb | Group | -0.229 | 0.107 | -0.441 | -0.019 | 1.003 | 1500 |
| R7 | O2Hb | Q-value | -0.070 | 0.078 | -0.221 | 0.084 | 1.000 | 5612 |
| R7 | O2Hb | Group x Q-value | 0.069 | 0.110 | -0.148 | 0.288 | 1.001 | 5852 |
| R7 | HHb | Intercept | 0.014 | 0.078 | -0.140 | 0.165 | 1.001 | 1331 |
| R7 | HHb | Group | 0.088 | 0.111 | -0.126 | 0.306 | 1.002 | 1549 |
| R7 | HHb | Q-value | -0.070 | 0.079 | -0.222 | 0.089 | 1.001 | 4904 |
| R7 | HHb | Group x Q-value | 0.007 | 0.112 | -0.217 | 0.222 | 1.000 | 4978 |
| R8 | O2Hb | Intercept | -0.017 | 0.086 | -0.184 | 0.154 | 1.004 | 675.9 |

| | | | | | | | | |
|---|---|---|---|---|---|---|---|---|
| R8 | O2Hb | Group | 0.152 | 0.125 | -0.089 | 0.401 | 1.009 | 649.0 |
| R8 | O2Hb | Q-value | -0.047 | 0.130 | -0.305 | 0.207 | 1.002 | 1337 |
| R8 | O2Hb | Group x Q-value | -0.131 | 0.187 | -0.495 | 0.242 | 1.008 | 1365 |
| R8 | HHb | Intercept | 0.025 | 0.082 | -0.133 | 0.189 | 1.002 | 923 |
| R8 | HHb | Group | 0.081 | 0.117 | -0.147 | 0.315 | 1.007 | 924.1 |
| R8 | HHb | Q-value | -0.197 | 0.183 | -0.561 | 0.156 | 1.001 | 1443 |
| R8 | HHb | Group x Q-value | 0.119 | 0.268 | -0.413 | 0.632 | 1.001 | 1279 |
| R9 | O2Hb | Intercept | 0.128 | 0.076 | -0.022 | 0.275 | 1.007 | 748.8 |
| R9 | O2Hb | Group | -0.236 | 0.105 | -0.441 | -0.032 | 1.003 | 739.7 |
| R9 | O2Hb | Q-value | -0.297 | 0.184 | -0.646 | 0.074 | 1.005 | 959.6 |
| R9 | O2Hb | Group x Q-value | 0.261 | 0.268 | -0.267 | 0.776 | 1.004 | 1075 |
| R9 | HHb | Intercept | 0.069 | 0.080 | -0.090 | 0.228 | 1.004 | 1431 |
| R9 | HHb | Group | -0.045 | 0.112 | -0.261 | 0.183 | 1.001 | 1613 |
| R9 | HHb | Q-value | -0.242 | 0.085 | -0.409 | -0.071 | 1.001 | 4948 |
| R9 | HHb | Group x Q-value | 0.191 | 0.121 | -0.049 | 0.425 | 1.000 | 4469 |
| R10 | O2Hb | Intercept | 0.021 | 0.065 | -0.105 | 0.149 | 1.002 | 1122 |
| R10 | O2Hb | Group | -0.014 | 0.093 | -0.199 | 0.167 | 1.002 | 1123 |
| R10 | O2Hb | Q-value | 0.007 | 0.071 | -0.133 | 0.147 | 1.001 | 3570 |
| R10 | O2Hb | Group x Q-value | -0.141 | 0.100 | -0.342 | 0.054 | 1.001 | 3278 |
| R10 | HHb | Intercept | 0.031 | 0.059 | -0.087 | 0.143 | 1.003 | 1340 |
| R10 | HHb | Group | 0.030 | 0.084 | -0.133 | 0.194 | 1.002 | 1579 |
| R10 | HHb | Q-value | -0.111 | 0.054 | -0.220 | -0.005 | 1.002 | 2185 |
| R10 | HHb | Group x Q-value | 0.079 | 0.078 | -0.075 | 0.233 | 1.000 | 2616 |

LMM – Linear mixed-effect model; O2Hb – Oxygenated hemoglobin; HHb – Deoxygenated hemoglobin; CI – Confidence interval; ESS – Effective sample size; s.e.m. – Standard error of mean; CT – Control subjects with no criminal record; TR-K –Theft recidivist without kleptomania.

**Supplementary Table S3. A summary of the Bayesian LMM results for associations of RPE with PFC responses.**

**Estimates (difference from Intercept for Group, Q-value, and Group x Q-value)**

| Site | Signal | Term | Level | Estimate (β) | s.e.m. | 95% CI | | $\hat{R}$ | ESS (bulk) | ESS (tail) |
|---|---|---|---|---|---|---|---|---|---|---|
| | | | | | | Lower | Upper | | | |
| R1 | O2Hb | Intercept | | 0.022 | 0.100 | -0.173 | 0.219 | 1.002 | 971.1 | 1934 |
| R1 | O2Hb | Group | TR-K | 0.135 | 0.090 | -0.041 | 0.318 | 1.003 | 935.7 | 1839 |
| R1 | O2Hb | Group | CT | -0.109 | 0.093 | -0.297 | 0.072 | 1.003 | 972.2 | 1775 |
| R1 | O2Hb | RPE | | 0.049 | 0.050 | -0.050 | 0.148 | 1.001 | 1456 | 3327 |
| R1 | O2Hb | Group x RPE | TR-K | -0.022 | 0.078 | -0.177 | 0.128 | 1.001 | 1575 | 3360 |
| R1 | O2Hb | Group x RPE | CT | 0.120 | 0.063 | -0.004 | 0.247 | 1.002 | 1245 | 2865 |
| R1 | HHb | Intercept | | -0.011 | 0.091 | -0.192 | 0.171 | 1.009 | 561.0 | 1252 |
| R1 | HHb | Group | TR-K | 0.091 | 0.085 | -0.073 | 0.249 | 1.003 | 616.7 | 1398 |
| R1 | HHb | Group | CT | -0.044 | 0.087 | -0.210 | 0.129 | 1.003 | 660.9 | 1438 |
| R1 | HHb | RPE | | 0.088 | 0.050 | -0.012 | 0.186 | 1.003 | 1558 | 4737 |
| R1 | HHb | Group x RPE | TR-K | 0.086 | 0.081 | -0.071 | 0.241 | 1.001 | 1837 | 5169 |
| R1 | HHb | Group x RPE | CT | 0.089 | 0.062 | -0.034 | 0.213 | 1.003 | 1279 | 3337 |
| R2 | O2Hb | Intercept | | 0.010 | 0.072 | -0.136 | 0.157 | 1.003 | 1318 | 2604 |
| R2 | O2Hb | Group | TR-K | 0.131 | 0.079 | -0.024 | 0.289 | 1.001 | 1288 | 2992 |
| R2 | O2Hb | Group | CT | -0.117 | 0.078 | -0.271 | 0.034 | 1.001 | 1268 | 3005 |
| R2 | O2Hb | RPE | | 0.025 | 0.058 | -0.086 | 0.138 | 1.000 | 8884 | 11580 |
| R2 | O2Hb | Group x RPE | TR-K | 0.058 | 0.091 | -0.118 | 0.235 | 1.000 | 9587 | 12190 |
| R2 | O2Hb | Group x RPE | CT | -0.008 | 0.070 | -0.149 | 0.128 | 1.000 | 9776 | 12440 |
| R2 | HHb | Intercept | | -0.019 | 0.074 | -0.172 | 0.130 | 1.005 | 1030 | 2026 |
| R2 | HHb | Group | TR-K | 0.049 | 0.087 | -0.120 | 0.217 | 1.002 | 936.3 | 1985 |
| R2 | HHb | Group | CT | 0.012 | 0.085 | -0.148 | 0.175 | 1.002 | 901.3 | 2054 |
| R2 | HHb | RPE | | 0.114 | 0.061 | -0.007 | 0.233 | 1.000 | 4849 | 8557 |
| R2 | HHb | Group x RPE | TR-K | 0.187 | 0.095 | 0.001 | 0.378 | 1.001 | 5144 | 9005 |
| R2 | HHb | Group x RPE | CT | 0.041 | 0.077 | -0.109 | 0.191 | 1.000 | 4130 | 9201 |
| R3 | O2Hb | Intercept | | -0.013 | 0.090 | -0.188 | 0.166 | 1.001 | 1272 | 1983 |
| R3 | O2Hb | Group | TR-K | 0.108 | 0.077 | -0.044 | 0.261 | 1.003 | 1292 | 2376 |
| R3 | O2Hb | Group | CT | -0.055 | 0.082 | -0.215 | 0.105 | 1.003 | 1422 | 2832 |
| R3 | O2Hb | RPE | | 0.099 | 0.072 | -0.046 | 0.243 | 1.001 | 2551 | 5529 |
| R3 | O2Hb | Group x RPE | TR-K | 0.139 | 0.113 | -0.089 | 0.360 | 1.002 | 3006 | 6240 |
| R3 | O2Hb | Group x RPE | CT | 0.060 | 0.088 | -0.121 | 0.236 | 1.001 | 2067 | 4142 |
| R3 | HHb | Intercept | | 0.038 | 0.096 | -0.149 | 0.226 | 1.005 | 848.7 | 1729 |
| R3 | HHb | Group | TR-K | 0.082 | 0.082 | -0.084 | 0.246 | 1.003 | 665.0 | 1610 |
| R3 | HHb | Group | CT | -0.112 | 0.088 | -0.283 | 0.059 | 1.003 | 787.3 | 1639 |
| R3 | HHb | RPE | | -0.056 | 0.100 | -0.252 | 0.145 | 1.003 | 1287 | 3274 |
| R3 | HHb | Group x RPE | TR-K | -0.176 | 0.156 | -0.487 | 0.134 | 1.002 | 1431 | 2878 |
| R3 | HHb | Group x RPE | CT | 0.064 | 0.131 | -0.190 | 0.317 | 1.002 | 1064 | 2320 |
| R4 | O2Hb | Intercept | | -0.019 | 0.086 | -0.192 | 0.159 | 1.003 | 1441 | 2565 |
| R4 | O2Hb | Group | TR-K | 0.105 | 0.082 | -0.056 | 0.264 | 1.003 | 1348 | 2957 |
| R4 | O2Hb | Group | CT | -0.028 | 0.084 | -0.193 | 0.135 | 1.002 | 1462 | 2965 |
| R4 | O2Hb | RPE | | 0.144 | 0.075 | -0.007 | 0.293 | 1.001 | 3760 | 6748 |
| R4 | O2Hb | Group x RPE | TR-K | 0.196 | 0.118 | -0.038 | 0.424 | 1.000 | 4096 | 7672 |
| R4 | O2Hb | Group x RPE | CT | 0.092 | 0.094 | -0.096 | 0.279 | 1.001 | 3376 | 7205 |
| R4 | HHb | Intercept | | -0.080 | 0.109 | -0.300 | 0.142 | 1.002 | 1049 | 2020 |
| R4 | HHb | Group | TR-K | 0.014 | 0.088 | -0.160 | 0.188 | 1.003 | 1179 | 2871 |
| R4 | HHb | Group | CT | 0.102 | 0.096 | -0.088 | 0.297 | 1.002 | 1631 | 3390 |
| R4 | HHb | RPE | | 0.216 | 0.145 | -0.077 | 0.509 | 1.001 | 1466 | 2812 |
| R4 | HHb | Group x RPE | TR-K | 0.500 | 0.229 | 0.049 | 0.960 | 1.001 | 1934 | 3418 |
| R4 | HHb | Group x RPE | CT | -0.069 | 0.183 | -0.429 | 0.296 | 1.002 | 1159 | 2687 |
| R5 | O2Hb | Intercept | | -0.005 | 0.080 | -0.164 | 0.152 | 1.007 | 1150 | 2565 |
| R5 | O2Hb | Group | TR-K | -0.013 | 0.088 | -0.185 | 0.161 | 1.002 | 1135 | 2083 |
| R5 | O2Hb | Group | CT | 0.018 | 0.087 | -0.153 | 0.188 | 1.002 | 1100 | 2311 |
| R5 | O2Hb | RPE | | 0.010 | 0.069 | -0.125 | 0.150 | 1.000 | 6325 | 8326 |
| R5 | O2Hb | Group x RPE | TR-K | 0.121 | 0.110 | -0.098 | 0.341 | 1.001 | 6154 | 8875 |
| R5 | O2Hb | Group x RPE | CT | -0.101 | 0.085 | -0.271 | 0.069 | 1.000 | 6559 | 9913 |
| R5 | HHb | Intercept | | 0.012 | 0.084 | -0.152 | 0.172 | 1.005 | 947.8 | 2434 |
| R5 | HHb | Group | TR-K | 0.089 | 0.082 | -0.068 | 0.249 | 1.003 | 1248 | 2535 |

| | | | | | | | | | | |
|---|---|---|---|---|---|---|---|---|---|---|
| R5 | HHb | Group | CT | -0.060 | 0.083 | -0.221 | 0.099 | 1.003 | 1280 | 2239 |
| R5 | HHb | RPE | | 0.052 | 0.066 | -0.076 | 0.184 | 1.001 | 5588 | 7945 |
| R5 | HHb | Group x RPE | TR-K | 0.110 | 0.104 | -0.087 | 0.315 | 1.001 | 5359 | 8332 |
| R5 | HHb | Group x RPE | CT | -0.005 | 0.081 | -0.165 | 0.154 | 1.000 | 4372 | 8212 |
| R6 | O2Hb | Intercept | | -0.044 | 0.088 | -0.230 | 0.134 | 1.003 | 757.3 | 1218 |
| R6 | O2Hb | Group | TR-K | 0.003 | 0.087 | -0.172 | 0.169 | 1.005 | 807.9 | 1856 |
| R6 | O2Hb | Group | CT | 0.059 | 0.089 | -0.118 | 0.234 | 1.003 | 818.7 | 1665 |
| R6 | O2Hb | RPE | | 0.116 | 0.081 | -0.042 | 0.279 | 1.002 | 2284 | 4898 |
| R6 | O2Hb | Group x RPE | TR-K | 0.149 | 0.127 | -0.101 | 0.399 | 1.000 | 2187 | 4274 |
| R6 | O2Hb | Group x RPE | CT | 0.084 | 0.102 | -0.119 | 0.290 | 1.003 | 1597 | 3821 |
| R6 | HHb | Intercept | | -0.046 | 0.090 | -0.226 | 0.131 | 1.002 | 1239 | 2649 |
| R6 | HHb | Group | TR-K | -0.018 | 0.089 | -0.196 | 0.152 | 1.002 | 1241 | 2227 |
| R6 | HHb | Group | CT | 0.080 | 0.088 | -0.094 | 0.258 | 1.002 | 1289 | 2644 |
| R6 | HHb | RPE | | 0.115 | 0.072 | -0.030 | 0.260 | 1.001 | 4660 | 8265 |
| R6 | HHb | Group x RPE | TR-K | 0.160 | 0.112 | -0.067 | 0.385 | 1.001 | 5291 | 9210 |
| R6 | HHb | Group x RPE | CT | 0.070 | 0.090 | -0.109 | 0.252 | 1.000 | 4343 | 7073 |
| R7 | O2Hb | Intercept | | 0.021 | 0.085 | -0.146 | 0.184 | 1.003 | 2049 | 3697 |
| R7 | O2Hb | Group | TR-K | 0.162 | 0.078 | 0.006 | 0.318 | 1.004 | 2038 | 4587 |
| R7 | O2Hb | Group | CT | -0.132 | 0.082 | -0.290 | 0.025 | 1.003 | 2155 | 4276 |
| R7 | O2Hb | RPE | | 0.055 | 0.069 | -0.085 | 0.195 | 1.000 | 6760 | 9998 |
| R7 | O2Hb | Group x RPE | TR-K | 0.136 | 0.108 | -0.080 | 0.357 | 1.000 | 7335 | 10370 |
| R7 | O2Hb | Group x RPE | CT | -0.026 | 0.086 | -0.201 | 0.148 | 1.001 | 5948 | 9426 |
| R7 | HHb | Intercept | | -0.023 | 0.083 | -0.194 | 0.138 | 1.001 | 2074 | 3071 |
| R7 | HHb | Group | TR-K | -0.042 | 0.079 | -0.201 | 0.113 | 1.001 | 2167 | 3704 |
| R7 | HHb | Group | CT | 0.079 | 0.080 | -0.079 | 0.238 | 1.001 | 2193 | 4242 |
| R7 | HHb | RPE | | 0.068 | 0.064 | -0.062 | 0.198 | 1.000 | 7506 | 10050 |
| R7 | HHb | Group x RPE | TR-K | 0.081 | 0.100 | -0.122 | 0.286 | 1.001 | 7569 | 9582 |
| R7 | HHb | Group x RPE | CT | 0.054 | 0.081 | -0.106 | 0.217 | 1.000 | 7257 | 10710 |
| R8 | O2Hb | Intercept | | -0.053 | 0.095 | -0.240 | 0.133 | 1.004 | 582.1 | 1485 |
| R8 | O2Hb | Group | TR-K | -0.078 | 0.089 | -0.251 | 0.090 | 1.004 | 643.6 | 1762 |
| R8 | O2Hb | Group | CT | 0.110 | 0.092 | -0.065 | 0.285 | 1.001 | 757.9 | 2046 |
| R8 | O2Hb | RPE | | 0.059 | 0.109 | -0.159 | 0.275 | 1.004 | 1209 | 3265 |
| R8 | O2Hb | Group x RPE | TR-K | -0.009 | 0.172 | -0.348 | 0.328 | 1.002 | 1440 | 3277 |
| R8 | O2Hb | Group x RPE | CT | 0.127 | 0.143 | -0.158 | 0.407 | 1.005 | 846 | 2361 |
| R8 | HHb | Intercept | | -0.066 | 0.107 | -0.278 | 0.147 | 1.002 | 1141 | 2793 |
| R8 | HHb | Group | TR-K | -0.026 | 0.089 | -0.199 | 0.146 | 1.001 | 1174 | 2127 |
| R8 | HHb | Group | CT | 0.114 | 0.094 | -0.069 | 0.305 | 1.001 | 1426 | 2944 |
| R8 | HHb | RPE | | 0.164 | 0.131 | -0.094 | 0.420 | 1.003 | 1420 | 3384 |
| R8 | HHb | Group x RPE | TR-K | 0.200 | 0.198 | -0.192 | 0.598 | 1.002 | 2135 | 4223 |
| R8 | HHb | Group x RPE | CT | 0.127 | 0.163 | -0.195 | 0.462 | 1.002 | 1158 | 2306 |
| R9 | O2Hb | Intercept | | -0.051 | 0.093 | -0.233 | 0.136 | 1.002 | 1127 | 2391 |
| R9 | O2Hb | Group | TR-K | 0.203 | 0.082 | 0.036 | 0.363 | 1.004 | 1074 | 2077 |
| R9 | O2Hb | Group | CT | -0.034 | 0.090 | -0.208 | 0.143 | 1.002 | 1127 | 2589 |
| R9 | O2Hb | RPE | | 0.314 | 0.119 | 0.070 | 0.547 | 1.004 | 1641 | 3306 |
| R9 | O2Hb | Group x RPE | TR-K | 0.544 | 0.187 | 0.166 | 0.911 | 1.002 | 1764 | 3616 |
| R9 | O2Hb | Group x RPE | CT | 0.085 | 0.149 | -0.204 | 0.377 | 1.005 | 1562 | 3249 |
| R9 | HHb | Intercept | | -0.026 | 0.080 | -0.186 | 0.128 | 1.002 | 764.2 | 1882 |
| R9 | HHb | Group | TR-K | 0.048 | 0.077 | -0.104 | 0.201 | 1.003 | 955.7 | 1857 |
| R9 | HHb | Group | CT | 0.026 | 0.078 | -0.127 | 0.181 | 1.003 | 972.3 | 1804 |
| R9 | HHb | RPE | | 0.138 | 0.064 | 0.016 | 0.263 | 1.000 | 5985 | 9849 |
| R9 | HHb | Group x RPE | TR-K | 0.240 | 0.098 | 0.048 | 0.436 | 1.001 | 6084 | 8864 |
| R9 | HHb | Group x RPE | CT | 0.036 | 0.078 | -0.115 | 0.191 | 1.001 | 5584 | 9595 |
| R10 | O2Hb | Intercept | | -0.010 | 0.079 | -0.167 | 0.149 | 1.003 | 1004 | 2017 |
| R10 | O2Hb | Group | TR-K | 0.061 | 0.076 | -0.082 | 0.210 | 1.003 | 1076 | 2388 |
| R10 | O2Hb | Group | CT | -0.011 | 0.077 | -0.168 | 0.139 | 1.005 | 1103 | 2391 |
| R10 | O2Hb | RPE | | 0.093 | 0.059 | -0.026 | 0.211 | 1.002 | 3266 | 7278 |
| R10 | O2Hb | Group x RPE | TR-K | 0.066 | 0.095 | -0.121 | 0.250 | 1.002 | 3731 | 8619 |
| R10 | O2Hb | Group x RPE | CT | 0.120 | 0.072 | -0.026 | 0.262 | 1.000 | 3353 | 7438 |
| R10 | HHb | Intercept | | -0.033 | 0.078 | -0.190 | 0.122 | 1.004 | 1336 | 2383 |
| R10 | HHb | Group | TR-K | -0.022 | 0.077 | -0.175 | 0.129 | 1.002 | 1694 | 2870 |
| R10 | HHb | Group | CT | 0.067 | 0.078 | -0.085 | 0.224 | 1.002 | 1717 | 2870 |
| R10 | HHb | RPE | | 0.085 | 0.036 | 0.015 | 0.154 | 1.001 | 13820 | 12120 |
| R10 | HHb | Group x RPE | TR-K | 0.098 | 0.057 | -0.014 | 0.210 | 1.000 | 13360 | 11420 |

| | | | | | | | | | | |
|---|---|---|---|---|---|---|---|---|---|---|
| R10 | HHb | Group x RPE | CT | 0.071 | 0.043 | -0.015 | 0.155 | 1.000 | 13890 | 11940 |

**Fixed Effects Estimates**

| Site | Signal | Term | Estimate (β) | s.e.m. | 95% CI | | $\hat{R}$ | ESS |
|---|---|---|---|---|---|---|---|---|
| | | | | | Lower | Upper | | |
| R1 | O2Hb | Intercept | 0.022 | 0.100 | -0.173 | 0.219 | 1.002 | 954.2 |
| R1 | O2Hb | Group | -0.199 | 0.145 | -0.489 | 0.081 | 1.003 | 944.3 |
| R1 | O2Hb | RPE | 0.049 | 0.050 | -0.050 | 0.148 | 1.001 | 1435 |
| R1 | O2Hb | Group x RPE | 0.101 | 0.072 | -0.039 | 0.243 | 1.002 | 1386 |
| R1 | HHb | Intercept | -0.011 | 0.092 | -0.192 | 0.171 | 1.009 | 562.8 |
| R1 | HHb | Group | -0.097 | 0.129 | -0.343 | 0.163 | 1.003 | 610.1 |
| R1 | HHb | RPE | 0.088 | 0.051 | -0.012 | 0.186 | 1.003 | 1562 |
| R1 | HHb | Group x RPE | 0.002 | 0.072 | -0.139 | 0.142 | 1.002 | 1611 |
| R2 | O2Hb | Intercept | 0.010 | 0.074 | -0.136 | 0.157 | 1.001 | 1316 |
| R2 | O2Hb | Group | -0.163 | 0.106 | -0.371 | 0.045 | 1.001 | 1238 |
| R2 | O2Hb | RPE | 0.025 | 0.057 | -0.086 | 0.138 | 1.000 | 8789 |
| R2 | O2Hb | Group x RPE | -0.047 | 0.081 | -0.204 | 0.111 | 1.000 | 9937 |
| R2 | HHb | Intercept | -0.019 | 0.076 | -0.172 | 0.130 | 1.005 | 1017 |
| R2 | HHb | Group | 0.001 | 0.110 | -0.211 | 0.217 | 1.002 | 886.5 |
| R2 | HHb | RPE | 0.114 | 0.062 | -0.007 | 0.233 | 1.000 | 4815 |
| R2 | HHb | Group x RPE | -0.103 | 0.087 | -0.275 | 0.068 | 1.000 | 4802 |
| R3 | O2Hb | Intercept | -0.013 | 0.091 | -0.188 | 0.166 | 1.001 | 1262 |
| R3 | O2Hb | Group | -0.100 | 0.125 | -0.347 | 0.147 | 1.003 | 1272 |
| R3 | O2Hb | RPE | 0.099 | 0.073 | -0.046 | 0.243 | 1.001 | 2537 |
| R3 | O2Hb | Group x RPE | -0.056 | 0.101 | -0.253 | 0.149 | 1.003 | 2541 |
| R3 | HHb | Intercept | 0.038 | 0.097 | -0.149 | 0.226 | 1.005 | 849.4 |
| R3 | HHb | Group | -0.182 | 0.137 | -0.453 | 0.085 | 1.002 | 656.6 |
| R3 | HHb | RPE | -0.056 | 0.101 | -0.252 | 0.145 | 1.003 | 1294 |
| R3 | HHb | Group x RPE | 0.170 | 0.147 | -0.116 | 0.460 | 1.001 | 1175 |
| R4 | O2Hb | Intercept | -0.019 | 0.089 | -0.192 | 0.159 | 1.003 | 1427 |
| R4 | O2Hb | Group | -0.074 | 0.125 | -0.320 | 0.171 | 1.003 | 1300 |
| R4 | O2Hb | RPE | 0.144 | 0.076 | -0.007 | 0.293 | 1.001 | 3731 |
| R4 | O2Hb | Group x RPE | -0.073 | 0.107 | -0.282 | 0.136 | 1.001 | 3767 |
| R4 | HHb | Intercept | -0.080 | 0.112 | -0.300 | 0.142 | 1.002 | 1028 |
| R4 | HHb | Group | 0.171 | 0.156 | -0.135 | 0.478 | 1.003 | 1243 |
| R4 | HHb | RPE | 0.216 | 0.149 | -0.077 | 0.509 | 1.001 | 1445 |
| R4 | HHb | Group x RPE | -0.402 | 0.208 | -0.814 | 0.004 | 1.002 | 1598 |
| R5 | O2Hb | Intercept | -0.005 | 0.081 | -0.164 | 0.152 | 1.007 | 1141 |
| R5 | O2Hb | Group | 0.064 | 0.121 | -0.178 | 0.297 | 1.002 | 1050 |
| R5 | O2Hb | RPE | 0.010 | 0.070 | -0.125 | 0.150 | 1.000 | 6299 |
| R5 | O2Hb | Group x RPE | -0.157 | 0.100 | -0.357 | 0.036 | 1.000 | 6289 |
| R5 | HHb | Intercept | 0.012 | 0.083 | -0.152 | 0.172 | 1.005 | 989.6 |
| R5 | HHb | Group | -0.083 | 0.118 | -0.310 | 0.146 | 1.003 | 1144 |
| R5 | HHb | RPE | 0.052 | 0.066 | -0.076 | 0.184 | 1.001 | 5573 |
| R5 | HHb | Group x RPE | -0.082 | 0.093 | -0.266 | 0.100 | 1.000 | 4219 |
| R6 | O2Hb | Intercept | -0.044 | 0.092 | -0.230 | 0.134 | 1.003 | 743.5 |
| R6 | O2Hb | Group | 0.052 | 0.129 | -0.200 | 0.307 | 1.005 | 704.6 |
| R6 | O2Hb | RPE | 0.116 | 0.082 | -0.042 | 0.279 | 1.002 | 2287 |
| R6 | O2Hb | Group x RPE | -0.046 | 0.116 | -0.273 | 0.187 | 1.001 | 1544 |
| R6 | HHb | Intercept | -0.046 | 0.091 | -0.226 | 0.131 | 1.001 | 1243 |
| R6 | HHb | Group | 0.087 | 0.128 | -0.162 | 0.340 | 1.001 | 1205 |
| R6 | HHb | RPE | 0.115 | 0.073 | -0.030 | 0.260 | 1.001 | 4661 |
| R6 | HHb | Group x RPE | -0.064 | 0.104 | -0.266 | 0.142 | 1.000 | 5079 |
| R7 | O2Hb | Intercept | 0.021 | 0.084 | -0.146 | 0.184 | 1.003 | 2043 |
| R7 | O2Hb | Group | -0.177 | 0.118 | -0.411 | 0.055 | 1.004 | 1963 |
| R7 | O2Hb | RPE | 0.055 | 0.071 | -0.085 | 0.195 | 1.000 | 6729 |
| R7 | O2Hb | Group x RPE | -0.114 | 0.099 | -0.312 | 0.081 | 1.001 | 6648 |
| R7 | HHb | Intercept | -0.023 | 0.084 | -0.194 | 0.138 | 1.000 | 2064 |
| R7 | HHb | Group | 0.091 | 0.116 | -0.136 | 0.319 | 1.001 | 2062 |
| R7 | HHb | RPE | 0.068 | 0.065 | -0.062 | 0.198 | 1.000 | 7516 |
| R7 | HHb | Group x RPE | -0.019 | 0.093 | -0.202 | 0.163 | 1.001 | 7387 |
| R8 | O2Hb | Intercept | -0.053 | 0.096 | -0.240 | 0.133 | 1.004 | 573.2 |

| | | | | | | | | |
|---|---|---|---|---|---|---|---|---|
| R8 | O2Hb | Group | 0.107 | 0.138 | -0.161 | 0.375 | 1.003 | 626.3 |
| R8 | O2Hb | RPE | 0.059 | 0.110 | -0.159 | 0.275 | 1.004 | 1193 |
| R8 | O2Hb | Group x RPE | 0.096 | 0.160 | -0.220 | 0.411 | 1.003 | 1015 |
| R8 | HHb | Intercept | -0.066 | 0.108 | -0.278 | 0.147 | 1.002 | 1127 |
| R8 | HHb | Group | 0.113 | 0.151 | -0.182 | 0.407 | 1.000 | 1223 |
| R8 | HHb | RPE | 0.164 | 0.131 | -0.094 | 0.420 | 1.003 | 1409 |
| R8 | HHb | Group x RPE | -0.052 | 0.183 | -0.411 | 0.308 | 1.001 | 1709 |
| R9 | O2Hb | Intercept | -0.051 | 0.094 | -0.233 | 0.136 | 1.002 | 1121 |
| R9 | O2Hb | Group | -0.081 | 0.132 | -0.333 | 0.184 | 1.003 | 1014 |
| R9 | O2Hb | RPE | 0.314 | 0.121 | 0.070 | 0.547 | 1.004 | 1628 |
| R9 | O2Hb | Group x RPE | -0.324 | 0.167 | -0.656 | 0.003 | 1.002 | 1706 |
| R9 | HHb | Intercept | -0.026 | 0.080 | -0.186 | 0.128 | 1.002 | 764.7 |
| R9 | HHb | Group | 0.023 | 0.110 | -0.191 | 0.238 | 1.004 | 907.1 |
| R9 | HHb | RPE | 0.138 | 0.064 | 0.016 | 0.263 | 1.000 | 5972 |
| R9 | HHb | Group x RPE | -0.144 | 0.088 | -0.319 | 0.030 | 1.002 | 5560 |
| R10 | O2Hb | Intercept | -0.010 | 0.080 | -0.167 | 0.149 | 1.003 | 993.6 |
| R10 | O2Hb | Group | -0.062 | 0.114 | -0.290 | 0.160 | 1.004 | 1044 |
| R10 | O2Hb | RPE | 0.093 | 0.060 | -0.026 | 0.211 | 1.002 | 3240 |
| R10 | O2Hb | Group x RPE | 0.039 | 0.084 | -0.125 | 0.204 | 1.001 | 3856 |
| R10 | HHb | Intercept | -0.033 | 0.079 | -0.190 | 0.122 | 1.004 | 1349 |
| R10 | HHb | Group | 0.068 | 0.113 | -0.152 | 0.289 | 1.002 | 1696 |
| R10 | HHb | RPE | 0.085 | 0.036 | 0.015 | 0.154 | 1.000 | 13660 |
| R10 | HHb | Group x RPE | -0.019 | 0.050 | -0.118 | 0.078 | 1.000 | 13240 |

LMM – Linear mixed-effect model; RPE – Reward prediction error; O2Hb – Oxygenated hemoglobin; HHb – Deoxygenated hemoglobin; CI – Confidence interval; ESS – Effective sample size; s.e.m. – Standard error of mean; CT – Control subjects with no criminal record; TR-K –Theft recidivist without kleptomania.

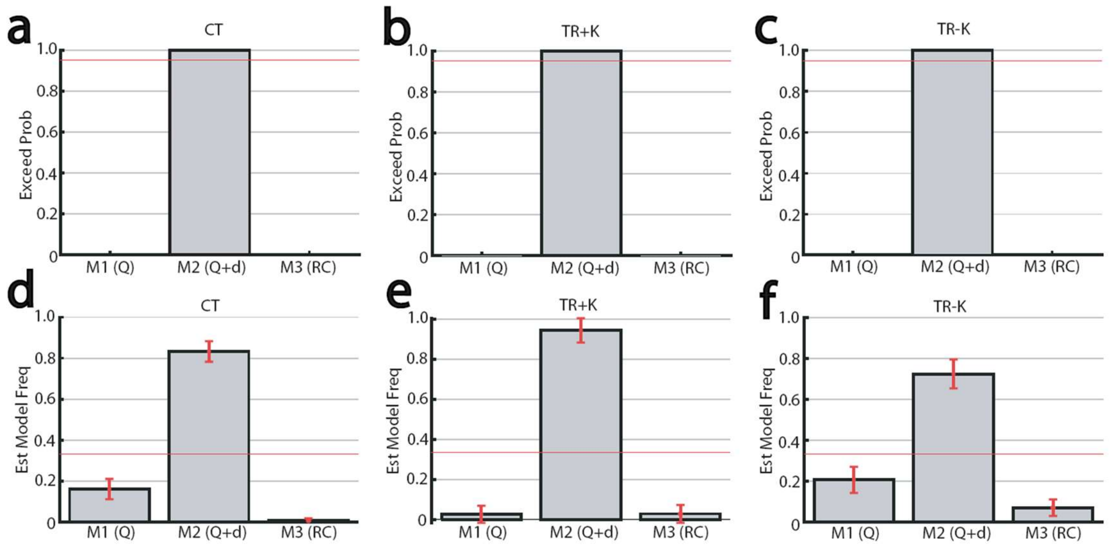


**Supplementary Figure S1. Group-level Bayesian Model Selection (BMS). a-c,** Graphs showing the exceedance probabilities for the standard Q-learning model (M1(Q)), the Q-learning model with decay (M2(Q+d)), and the random choice model (M3(RC)) in CT (a), TR+K, and TR-K (b), respectively. **d-f,** Graphs similar to a-c, but showing estimated model selection frequencies in CT (d), TR+K (e), and TR-K (f).

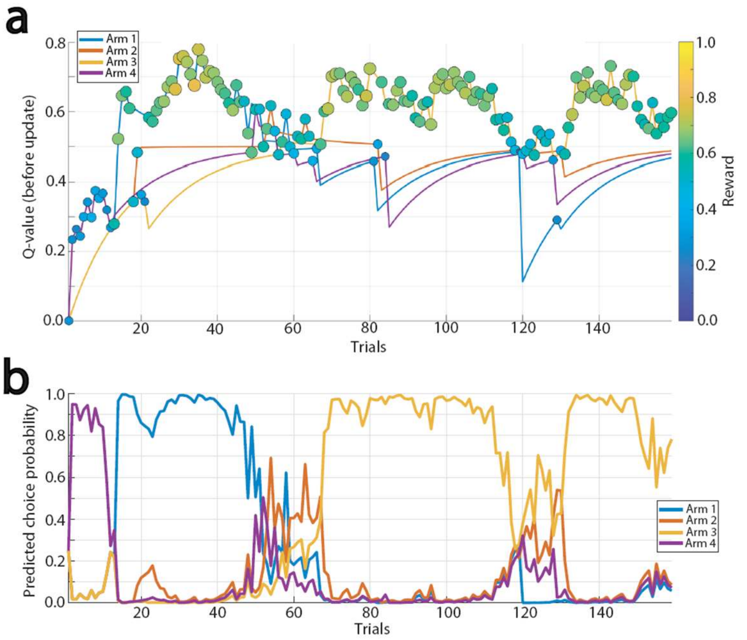


**Supplementary Figure S2. An example of the model evaluation of a single participant's choices. a,** A graph showing the model's internal Q-value (a) for each of the 4 arms progressing across the trials in a single represented participant. Four different colored lines represent each arm. The circles indicate which arm the participant chose on that specific trial, illustrating when the participant wins, the Q-value line for that arm jumps up, and the participant's actual choice markers cluster around the arm with the highest Q-value. The negative log-likelihood (NLL) that measures the predictive error of the model is 81.61, and the accuracy is 86.20% in this example. Since the chance-level NLL for 160 trials is approximately 221.76, this indicates that the performance of the model is good and realistic. The NLL of 81.61 with 160 trials yields the average probability that the model assigned to the participant's actual choices is 60.05%. Given that in a 4-arm task, a random guess gives a 25% probability, the average probability over 60% across 160 trials indicates a high inverse temperature parameter (β) paired with occasional exploratory noise. **b,** A graph similar to a, but showing the predicted choice probability.

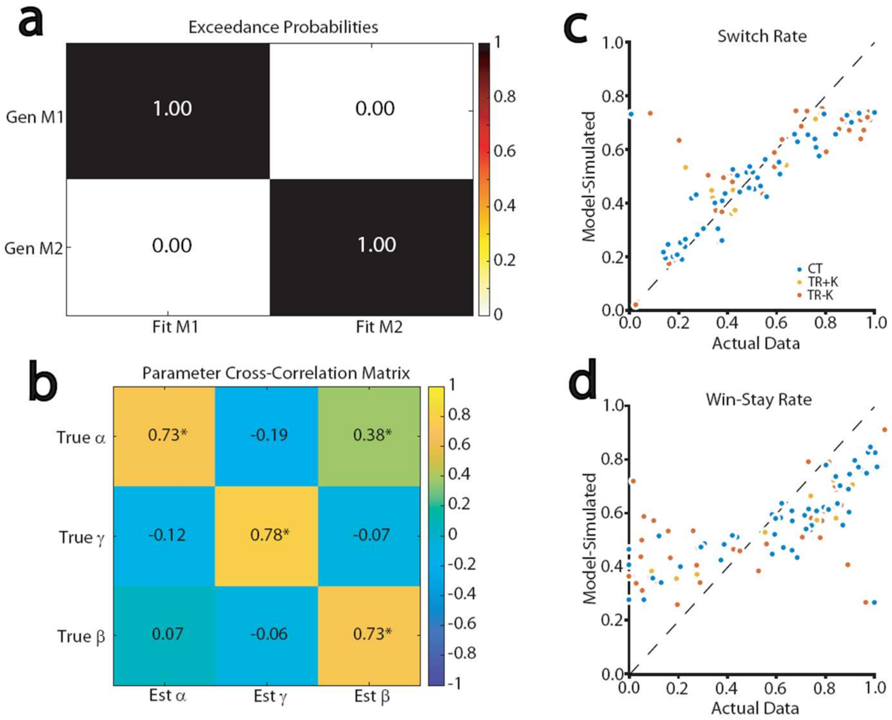


**Supplementary Figure S3. Validations of the model with model and parameter recovery analyses and behavioral reproduction. a,** A confusion matrix with the model recovery analysis, showing that the models were identifiable and not subject to mimicry. The analysis was conducted with the synthetic choice data for 25 agents per model using the generative parameters and task structure of the 160-trial 4-arm bandit task, along with cross-fitting for the standard Q-learning and Q-learning with decay models to create the matrix. **b,** A correlation matrix with the parameter recovery analysis for the Q-learning model with decay, showing that Person's correlation coefficients r, along with the color-coded heat map, between the generative parameters and the recovered parameters (α, β, and γ). A statistically significant ($p < 0.05$) correlation is indicated as an asterisk. The matrix was created with the100 synthetic dataset that successfully recovered the generative model. While parameter recovery correlations in extended tasks frequently exceed r = 0.80, the moderate correlations observed here were a consequence of the relatively brief task length (160 trials), which was a necessary methodological constraint. Nonetheless, the highly significant recovery correlations confirm that the parameters remain robustly identifiable and capture distinct mechanistic variance, thereby validating their use for group-level behavioral inferences. As is also frequently observed in reinforcement learning models, we noted a moderate positive cross-correlation between true learning rate and estimated inverse temperature (r = 0.38), reflecting the known computational trade-off between value-updating magnitude and choice determinism in shorter tasks. Crucially, the primary recovery correlations on the diagonal remained strictly stronger than these off-diagonal trade-offs, confirming that the model captures distinct, identifiable variance for each parameter. **c-d,** Scatter plots showing correlations between the simulated (with the Q-learning model with decay) and actual (empirically obtained) switch rate (c) and win-stay rate (d). While the model captured the overall variance and group-level differences effectively (yielding strong correlations across both metrics), minor deviations from the identity line were observed at the distribution extremes (ceilings and floors), which is a common artifact of bounded behavioral proportions and saturating Softmax functions.

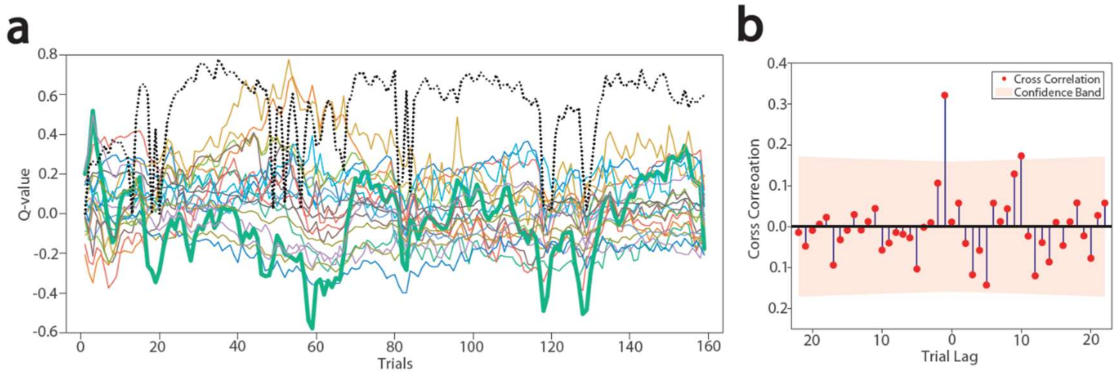


**Supplementary Figure S4. A single subject example of fNIRS signals with model-derived latent reinforcement variables. a,** A graph showing the trial-by-trial, standardized O2Hb and HHb signals across 10 sampling locations in the PFC and model's internal Q-value in a representative participant. A black dot line indicates the Q-value, and the solid color lines indicate each of the O2Hb and HHb signals in each location, with one of them highlighted by a bold line. **b,** A graph showing the cross-correlation analysis with differencing of the data between the Q-value and one of the fNIRS signals highlighted in a. A presence of the high correlation around 0 indicates near synchronous correlation between them.
